# Blocking Compensatory Matrix Cross-Linking Accelerates Thoracic Aortopathy in a Mouse Model of Marfan Syndrome

**DOI:** 10.64898/2026.08.24.746812

**Authors:** Gavin Mays, Jay D. Humphrey

**Affiliations:** Department of Biomedical Engineering Yale University, New Haven, CT, USA; Vascular Biology and Therapeutics Program Yale School of Medicine, New Haven, CT, USA

**Keywords:** fibrillin-1, lysyl oxidase, hypertension, homeostasis, aneurysm, dissection

## Abstract

Mechanical homeostasis plays a central role in promoting and preserving optimal structure and function in the adult aorta. Although pathogenic variants can compromise homeostatic processes, it appears that intramural cells yet attempt to compensate for some genetically induced changes. In particular, lysyl oxidase is higher in the adult Marfan aorta compared with the age-matched control aorta. Here, we block lysyl oxidase in adult *Fbn1^C1041G/+^* Marfan syndrome mice after stimulating aortic disease progression via induced hypertension. Whereas hypertension alone increases aortic dilatation, concurrent blocking of lysyl oxidase results in a dramatic increase in disease severity, driving an otherwise mild aortic phenotype in adult male *Fbn1^C1041G/+^* Marfan mice to aneurysmal dilatations as well as dissection and rupture, with frequent premature death. Deposition and cross-linking of fibrillar collagens, among other extracellular matrix constituents, can represent a protective compensation against severe disease in the Marfan aorta. The present study emphasizes the need clinically to avoid compromising new collagen deposition and suggests that strategies to augment collagen cross-linking could be beneficial.

## INTRODUCTION

Many pathogenic variants predispose to thoracic aortic disease, particularly variants in genes that encode key extracellular matrix (ECM) constituents that determine wall stiffness and strength and key intracellular components that dictate smooth muscle cell (SMC) contraction and relaxation (Brownstein et al. 2017; Milewicz et al. 2017). Pathogenic variants leading to aortopathy include, for example, those that disrupt the extracellular glycoprotein fibrillin-1 as well as those that disrupt the intracellular protein smooth muscle myosin heavy chain. Because these and similarly affected constituents fall along the mechanotransduction axis, compromised mechanosensing (assessment) and mechanoregulation (assembly) of ECM are thought to be particularly pathologic (Humphrey et al. 2015; Sengle and Sakai 2015; Ramirez et al. 2018; Yamashiro and Yanagisawa 2020; Creamer et al. 2021). Whereas it is axiomatic that pathogenic variants ultimately disrupt normal aortic structure and function, it appears prudent to pursue therapeutic options that both prevent pathologic consequences and promote protective compensations (Humphrey and Milewicz 2026). Toward this end, there is first a need to understand better the potential offsetting effects of compensatory adaptations.

Marfan syndrome (MFS) stems from pathogenic variants in the gene (*FBN1* in humans, *Fbn1* in mice) that encodes fibrillin-1, which normally contributes both to the long-term stability of the elastic fibers that endow the wall with distensibility and resilience and to the ability of SMCs to mechanosense their local mechanical environment (Van Andel et al. 2021; J. D. Humphrey and Tellides 2019). It has been found, however, that lysyl oxidase (LOX) is increased in the Marfan aorta (Busnadiego, Gorbenko del Blanco, et al. 2015), which appears to provide a compensatory protective structural benefit, at least in the *Fbn1^C1041G/+^* mouse model of MFS (Weiss et al. 2023). LOX is a key cross-linker of constituents within the ECM, particularly newly synthesized elastic fibers and fibrillar collagens. It appears that increasing the cross-linking of these constituents above normal can slow progressive dilatation (by delaying loss of elastic fibers) and reduce the risk of dissection or rupture (by strengthening collagen fibers) of the Marfan aorta. Conversely, pathogenic variants in the gene (*LOX*) that encodes this enzyme predispose to thoracic aortic disease in humans (Guo et al. 2016; Lee et al. 2016) while studies using *Lox^-/-^* mice confirm severe aortic disease with rupture during the perinatal period (Mäki et al. 2002; Kim et al. 2017; Staiculescu et al. 2017). It has also been shown that β-aminopropionitrile (BAPN) – an inhibitor of LOX and perhaps LOX-like family members – accelerates thoracic aortopathy in murine models (Zheng et al. 2020; Sawada et al. 2022), particularly in young animals wherein rates of synthesis of elastin and especially collagen are high (Kelleher et al. 2004; Weiss et al. 2024).

Hypertension is a key risk factor for thoracic aortic disease, including in MFS (Milewicz et al. 2021). Sustained elevations in blood pressure often increase turn-over of ECM constituents, such as collagen (Nissen et al. 1978), and thus induce marked changes within the ECM that alter biomechanical function of both the aortic media and adventitia (Bersi et al. 2017; Spronck et al. 2021). We previously showed computationally (Latorre and Humphrey 2020) and experimentally (Mays and Humphrey 2026) that hypertension can exacerbate aortic disease in cases of compromised elastic fiber integrity, as in MFS. We hypothesized here that blocking LOX activity during a hypertensive stimulus would increase disease severity in adult *Fbn1^C1041G/+^* mice, as evidenced by marked reductions in biomechanical function and structural integrity. The findings presented below support this hypothesis, confirming the critical importance of ECM cross-linking, particularly during periods of heightened ECM turnover in response to an external stressor such as hypertension.

## METHODS

### Mice

All procedures involving live mice were approved by the Institutional Animal Care and Use Committee (IACUC) of Yale University. To avoid conflating effects of natural postnatal development, all mice entered the study at 8 weeks of age, the previously identified age of biomechanical maturity of healthy male C57BL/6J thoracic aortas (Murtada et al. 2021). To examine sex and genotype as biological variables, we studied littermate male (M) and female (F) *Fbn1^+/+^*(WT) and *Fbn1^C1041G/+^* (MFS) mice. The *Fbn1^C1041G/+^* breeders were obtained from Jackson Laboratories (Bar Harbor, ME) and were on a C57BL/6J background. In accordance with ARRIVE guidelines, mice were randomly allocated to three basic study groups: normal-diet normotensive controls (NT), BAPN in the drinking water (BAPN), and hypertension-inducing chow and drinking water supplement + BAPN (HT+BAPN). The BAPN (β-aminopropionitrile; Sigma-Aldritch) was provided via the drinking water at 2.0 g/L. Blood pressure was elevated using 8% NaCl high-salt chow (Harland TD.92012) plus 3.0 g/L of L-NAME (N-nitro-L-arginine methyl ester; Thermo Scientific) in the drinking water, which was replaced thrice weekly. Examination across both sexes, two genotypes, and three diets yielded 12 age-matched experimental groups. Blood pressure was measured using a CODA tail-cuff device (Kent Scientific) at 12 weeks of age, the study endpoint, with a preceding three-day acclimation period to blood pressure measurement. Finally, mice were euthanized via CO_2_ inhalation followed by exsanguination upon removal of the thoracic aorta for biaxial testing. Any mice that died prior to the 12-week endpoint were necropsied (except in cases of cannibalism) to assess possible thoracic aortopathy but excluded from biomechanical phenotyping and histological examination.

### Biaxial Vessel Testing

Biomechanical phenotyping was limited to mice that survived to 12 weeks of age, hence introducing an unavoidable survivorship bias. To promote reproducibility, rigor, and robust comparison to other studies performed in our lab, we used previously validated experimental protocols, systems, and analyses to characterize the WT and MFS aortas (Cavinato et al. 2021; Weiss et al. 2023; Mays and Humphrey 2026). Briefly, the ascending aorta was excised from the aortic valve to the distal end of the aortic arch and cleaned of perivascular tissue. The vessel segment was gently perfused with a Hanks Buffered Saline Solution (HBSS) to visualize intramural blood, if any. Ligatures of unbraided 9-0 nylon sutures were used to close the base of the left common carotid and immediate distal aortic arch. The vessel was then cannulated on custom drawn glass cannulae through the aortic root (proximal end) and brachiocephalic artery (distal end) and secured using 6-0 silk sutures. Care was taken to not over-stretch or otherwise deform the vessel throughout this process. The cannulated vessel assembly was then placed within a custom computer-controlled biaxial testing device (Gleason et al. 2004) and immersed in an oxygenated (95% O_2_, 5% CO_2_) and warmed (37°C) Krebs-Ringer bicarbonate buffered solution (KREBS) containing 2.5 mM CaCl_2_.

Vasoactive testing was used to examine aortic responses. Each vessel was axially stretched to the specimen-specific, energetically preferred *in vivo* value 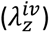 and held at a constant distending pressure (90 mmHg). Following a standardized *ex vivo* acclimation period, the isometrically and isobarically loaded vessel was vasoconstricted with 100 mM potassium chloride (KCl), vasorelaxed with a KREBS wash-out, vasoconstricted with 1 µM Angiotensin II (Ang-II), vasorelaxed with KREBS wash-out, vasoconstricted with 1 µM phenylephrine (PE; an alpha-1 adrenergic receptor activator in SMCs), vasodilated with 10 µM acetylcholine (Ach; an endothelial cell dependent stimulus of nitric oxide), and finally vasoconstricted with 1 mM L-NAME (an inhibitor of endothelial cell nitric oxide synthase). This active testing was completed within 4 hours of tissue extraction to ensure cell viability.

Passive testing was then performed to examine mechanical contributions of ECM constituents while rendering SMCs inactive. First, the KREBS solution was replaced with HBSS at room temperature to minimize smooth muscle cell contractility. While held at the energetically preferred axial stretch 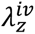, vessels were pre-conditioned by cyclic pressurization from 10-140 mmHg, then subjected to a sequence of seven computer-controlled biaxial protocols: three cyclic pressurization tests from 10-140 mmHg at three fixed axial stretches 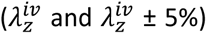 followed by four cyclic axial loading tests from 0 mN to a specimen specific *f_max_*(corresponding to force estimated at stretch 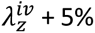 and distending pressure 140 mmHg) at four fixed luminal pressures (10, 60, 100, and 140 mmHg). Following passive testing, optical coherence tomography (OCT), enabled via a CALLISTO Spectral Domain OCT Imaging System (ThorLabs, Germany), was used to assess possible intramural voids, indicating spontaneous delamination or prior dissection. Vessels that did not undergo active testing were then committed to subsequent passive pressure-burst tests. Note that all tests neglect the mechanical contribution of perivascular tissue that may support against circumferential distension *in vivo* (Ferruzzi et al. 2018), especially in the case of highly diseased vessels in which perivascular tissue may be closely adherent to the vessel. Distending pressure, outer diameter, axial length and axial force measurements were recorded on-line throughout active and passive biaxial testing.

Pressure-burst tests examined mechanical properties up to a maximum distending pressure at rupture. Following the passive testing sequence, vessels were stretched to the aforementioned specimen-specific stretch 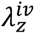 and pressurized briefly to 100 mmHg to verify no prior leakage. Then, vessels were pressurized monotonically from 10 mmHg until rupture. Luminal distending pressure and outer diameter were recorded on-line. For consistency, all samples were pressurized using HBSS at the same inlet flow rate. Following rupture, vessels were imaged to examine the gross location (both axially and circumferentially), orientation, and length of the rupture site. Using incompressibility assumptions, the bulk rupture stress was computed as the radially-averaged circumferential Cauchy stress at failure (calculated using *σ_θ_* = *P_a_*/ℎ wherein *P* is the distending pressure, *α* is the luminal radius, and ℎ is the wall thickness; Kawamura et al. 2021).

### Passive Property Quantification

We used methods shown previously to quantify well the passive mechanical behavior (Cavinato, Chen, et al. 2021; Dar Weiss et al. 2023), including pressure-diameter relationships, stress-stretch relationships, radially-averaged elastic stored energy, and biaxial stretch, stress, and stiffness. To determine these metrics, data from the unloading portion of each of the seven passive biaxial protocols (∼2800 data points) were fit simultaneously using an independently validated four-fiber family constitutive model:

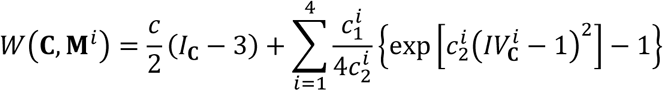

wherein *c*(kPa), 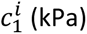, and 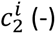 are free model parameters determined using a Levenberg-Marquardt nonlinear regression, with *i* = 1,2,3,4 representing the *i*^th^ family of fibers (with 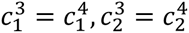 enforcing symmetric diagonal fibers). 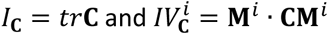 are coordinate invariant measures of deformation, with **C** = **F**^T^**F** the right Cauchy-Green tensor and **F** the deformation gradient tensor (*det* **F** = 1 because of assumed incompressibility), with superscript T denoting transpose. The *i*^th^ fiber family direction is given by unit vector 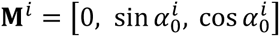, wherein axial 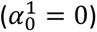, circumferential 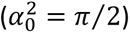, and two symmetric diagonal families of fibers 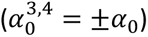 capture phenomenologically the complex biaxial material behavior that results from oriented structural fibers and cross-linking between. Biaxial material stiffness was computed from *W* using the theory of small deformations superimposed on large to calculate physiologically relevant values that are appropriate for simulations of fluid-solid-interactions (Baek et al. 2007) or fluid-solid-growth simulations (Schwarz et al. 2023). Overall distensibility was computed by 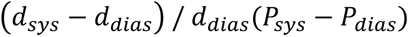 where *d* and *P* are diameter and pressure at systolic and diastolic values. Local pulse wave velocity was computed by the Moens-Korteweg equation (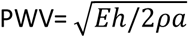, where *E* is the linearized circumferential material stiffness, ℎ wall thickness, *α* luminal radius, and *ρ* the mass density of the fluid). Pressure-dependent biaxial data metrics were computed at group-specific systolic blood pressure as well as at a fixed common pressure of 100 mmHg.

### Histology

Following active + passive mechanical testing, aortas were removed from the testing apparatus and fixed in their unloaded state in a 10% formalin solution for 24 hours before being transferred to a 70% ethanol solution for storage at 4°C. Samples selected at random for histological examination were embedded in paraffin, sectioned at 5-µm thickness, mounted on glass slides, and stained with Movat’s pentachrome stain (MOV), picrosirius red (PSR), or immunohistochemical stains (IHC). MOV was used to distinguish elastin and nuclei (black), collagen fibers (yellow), ground substance and glycosaminoglycans (blue), cell cytoplasm (light red) and fibrin (dark red). PSR enabled examination of thick (red/orange) and thin (green/yellow) collagen fibers. IHC stains included CD45 (pan leukocyte marker) and CD68 (lysosomal activity, often high in macrophages). Both MOV and IHC slides were imaged in brightfield mode. PSR slides were imaged under circularly polarized light microscopy with separate exposures to capture medial and adventitial areas (Cavinato, Murtada, et al. 2021). Images were acquired using an Olympus BX/51 microscope equipped with a DP70 digital camera and 20x objective (NA 0.50). When axial cross-sections exceeded the field-of-view, multiple images were acquired and stitched together using Image Composite Editor software (Microsoft Research). Based on prior studies, *n* = 4-5 randomly selected biological replicates were studied per group, each with *n* = 3 technical replicate sections per stain, resulting in *n* > 12 samples per group. To promote reproducibility, all embedding, sectioning, and staining were performed by Yale’s Pathology Tissue Services while imaging and analysis were conducted in our laboratory.

Custom software was used to quantify percent constituents in MOV sections, fiber area fractions in PSR sections, and IHC-positive areas in IHC images (https://github.com/yale-humphrey-lab/histology-analysis). Total medial and adventitial wall areas were obtained from MOV and PSR sections, with separate quantification of layer-specific areas eliminating possible loss of adventitial area in MOV sections due to background thresholding and loss of medial area in PSR sections due to unstained, non-collagenous material. Adjacent 5-µm-thick cross-sections were used between MOV and PSR to promote consistency across stains. With consideration to batch size limitations, samples were stained in batches with respect to sex. To accommodate possible batch-to-batch variability in staining between sexes, histological findings were considered in terms of fold- or percent-changes rather than absolute values. Non-parametric Spearman’s rank correlations were used to examine sample-specific relationships between histological findings and passive mechanical behavior (as computed from **Supplemental Data File S1**).

### Statistics

Data are shown in Figures as mean ± standard error of the mean (SEM) but tabulated as mean ± standard deviation (SD) unless otherwise specified. We provided both representations of data to describe the reliability of mean results (SEM) and variability within data (SD). Statistical analyses were performed using a standard one-way ANOVA with Tukey’s multiple comparisons test across all 12 groups for all vasoactive, passive, pressure-burst, and histological data. Non-parametric Spearman’s rank correlations were used to examine sample-specific relationships among histological findings and key vasoactive and passive mechanics. For all reported comparisons, a *p* < 0.05 was considered statistically significant with *p* values represented in figures as follows: * for *p* < 0.05, ** for *p* < 0.005, *** for *p* < 0.0005, and **** for *p* < 0.0001.

## RESULTS

### Mice

The original 256 study mice were distributed across 12 groups (**Flowchart S1**), with 193 (75%) surviving to the 12-week endpoint and yielding 171 (67% yield) ascending aortas for biomechanical and histological analyses (noting that 22 vessels were excluded from analysis due to mechanical damage. The highest premature mortality in any group (51 deaths out of 76 mice, or 67%) was experienced by the male MFS mice exposed to HT+BAPN (**Figure 1a, b**). Necropsy of these mice attributed 13 deaths to rupture of the thoracic aorta, with other deaths due to abdominal aortic rupture (3), veterinary-mandated euthanasia for humane concerns (21), or unknown causes (14). Other male mice experienced lower mortality, namely, 0% WT that were NT, 5% (1 of 20) WT with BAPN, 22% (5 of 23) WT with HT+BAPN, 0% MFS that were NT, and 6% (1 of 17) MFS with BAPN. Female mice experienced fewer premature deaths (**Supplemental Figure S1a**), though with ∼27% mortality (4 of 15) in female MFS mice with HT+BAPN. There was also ∼8% (1 of 13) mortality in female WT mice with HT+BAPN, but no deaths in the other four female groups. As with most studies of acute and chronic aortopathy, premature deaths introduced an inherent survivorship bias in which the most severe phenotypes were not available for testing.

**Figure 1.**
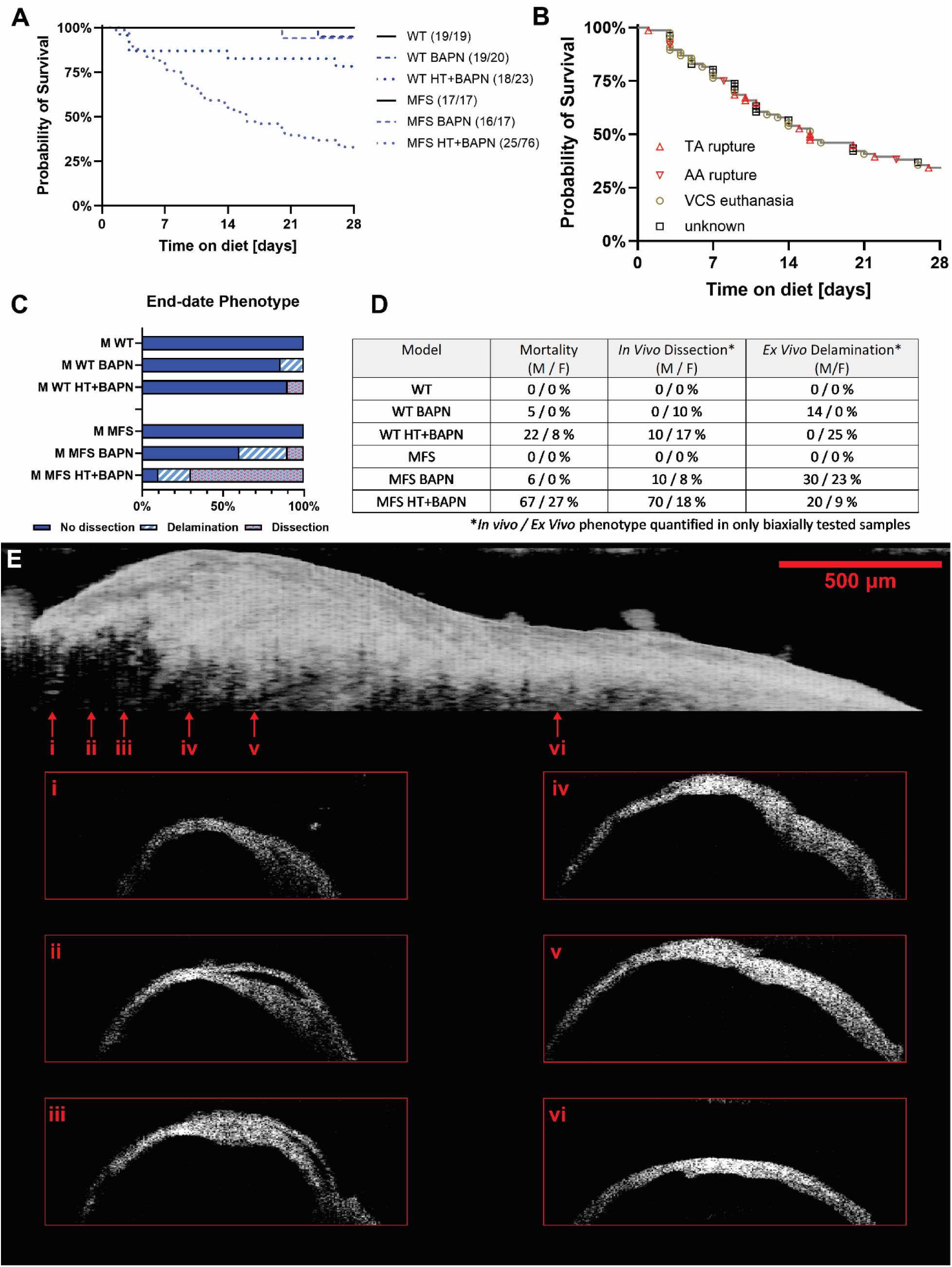
Male MFS mice exhibited the greatest early mortality and development of aortopathy. Male mice experienced greater early mortality than corresponding female mice under both BAPN and combined HT+BAPN challenge, with the greatest early mortality occurring in male MFS mice with combined HT+BAPN challenge (Panel A). Panel B indicates causes of death within the male MFS HT+BAPN group, by thoracic (TA) or abdominal (AA) aortic rupture, veterinary clinical service humane intervention (VCS), or unknown causes (most often unable to evaluate due to cannibalism). In particular, the male MFS HT+BAPN mice randomly selected for biaxial testing exhibited the highest rate of complex aortopathies, including 20% *ex vivo* delamination during biaxial testing and 70% *in vivo* dissection (Panels C, D). Phenotypes were confirmed following passive testing using optical coherence tomography, with Panel E showing the side profile from the outer curvature of an example dissected male MFS HT+BAPN sample with the aortic root oriented on the left. Note here that an intramural void is clearly seen starting on the outer curvature of the ascending aorta beginning near the aortic root (i) and extending approximately 500 µm axially (ii, iii) before termination (iv). Further along the length, while no voids may be present, there are clearly irregular areas of wall thickness, with local vulnerabilities both on the outer surface of the adventitia (v) and on the inner surface of the lumen (vi), which may act as stress foci in the case of aortic rupture. OCT image scale bar represents 500 µm with the six axial cross-sections represented by a 2000 µm x 700 µm rectangle.

Following passive biaxial testing (see below), OCT was used to assess the possible presence of two distinct types of intramural damage: a dissection defined by blood within the wall versus a spontaneous delamination in which *ex vivo* mechanical loading likely propagated previously imperceptible defects within the vessel wall (**Figure 1c-e**, **Supplemental Figure S1b**). The surviving male MFS mice with HT+BAPN again exhibited the most striking phenotypes, with a 70% (7 of 10) incidence of dissection and 20% (2 in 10) delamination amongst the biaxially tested vessels. In three cases, the vessels with *in vivo* dissections showed evidence of further *ex vivo* delamination. That is, only 10% of the aortas from the surviving 12-week-old male MFS mice exposed to HT+BAPN exhibited intact aortic walls at the end of mechanical testing. By contrast, the male WT HT+BAPN mice had only a 10% (1 in 10) incidence of *in vivo* dissection, with vessels otherwise appearing intact after *ex vivo* testing. BAPN alone was a moderate inducer of aortic vulnerability in male mice, with 30% (3 of 10) of the MFS aortas and ∼14% (2 of 14) of the WT aortas delaminating during testing and 10% (1 of 10) of the MFS aortas experiencing *in vivo* dissection. Note, therefore, that only the male groups that experienced premature death also showed evidence of aortic dissections or delamination in the surviving mice at the 12-week endpoint. Female mice presented a wider range of damage phenotypes, with HT+BAPN driving dissections or delaminations in both MFS (∼18% dissection, ∼9% delamination) and WT (∼17% dissection; 25% delamination) aortas. BAPN alone drove aortic dissection or delamination in female MFS mice (∼8% dissection, ∼23% delamination) but only a single case of dissection (representing 10%) in female WT mice. Interestingly, despite these vulnerabilities in the female Marfan aortas with BAPN, there were no associated premature deaths. Because of this strong sexual dimorphism, we focus primarily on data for the 6 groups of male mice, though data are compared for all 12 groups either in the main body or Supplemental Materials.

HT+BAPN induced a gradual increase in blood pressure in male mice (**Supplemental Figure S2a-b**), significantly increasing systolic blood pressure by ∼20% (from 121 to 145 mmHg) in WT mice but less so (∼3%, from 126 to 130 mmHg) in MFS mice. Female mice also experienced increased systolic blood pressure (**Supplemental Figure S2d-e**), albeit to similar degrees, with WT increased by ∼14% (122 to 139 mmHg) and MFS increased by ∼15% (120 to 138 mmHg). Interestingly, BAPN alone moderately increased systolic blood pressure of males (∼13% in WT, ∼6% in MFS) while having negligible effect on blood pressure in females (0% in WT, ∼3% in MFS). Mouse body mass was also significantly impacted by HT+BAPN, inducing a 20-28% reduction across all groups (**Supplemental Figure S2c, f**). BAPN alone induced a moderate 8-9% reduction in body mass across all groups except male MFS mice, in which body mass increased by ∼2%, suggesting an aversion to the hypertension-inducing high salt chow or L-NAME in the drinking water.

### Vasoactive Responses

The Marfan aorta exhibited compromised vasoactivity compared to the WT aorta in both NT males and females (**Figure 2, Supplemental Figures S3-S5, Supplemental Table S1**). Male WT aortas retained considerable vasoactive function in both the BAPN and HT+BAPN groups. In response to PE, vasoconstriction dropped only modestly for BAPN (from 30.1% to 24.3% reductions in outer diameter) while remaining nearly preserved for HT+BAPN (30.1% vs. 29.1%) (**Figure 2a, c**); in response to Ach, vasodilatation was yet diminished for both BAPN (40.1% to 35.2%) and HT+BAPN (40.1% to 22.0%) (**Figure 2d, f**). Male Marfan aortas experienced further diminished vasoactive function for BAPN (PE from 18.1% to 5.9%; Ach from 23.8% to 8.0%) (**Figure 2b, c**) and were rendered essentially inactive for HT+BAPN (PE from 18.1% to 1.1%; Ach from 23.8% to 0.3%) (**Figure 2e, f**). Similar results were found for alternative stimulants of SMCs and endothelial activity (**Supplemental Figure S3**). In response to KCl, SMC function was relatively preserved across all male WT groups while it was markedly lower in male MFS groups. In response to L-NAME *ex vivo*, male MFS groups had notably diminished vasoconstrictive response, while in WT groups, hypertension reduced vasoconstriction. Note that endothelial cell function may be considered preserved when vasodilation in response to Ach recovers or exceeds the vasoconstriction induced by preceding PE. These data suggest that, while overall vasoactive function reduced in male aortas for BAPN alone, HT+BAPN reduced endothelial function further, likely due to the prolonged delivery of L-NAME in drinking water.

**Figure 2.**
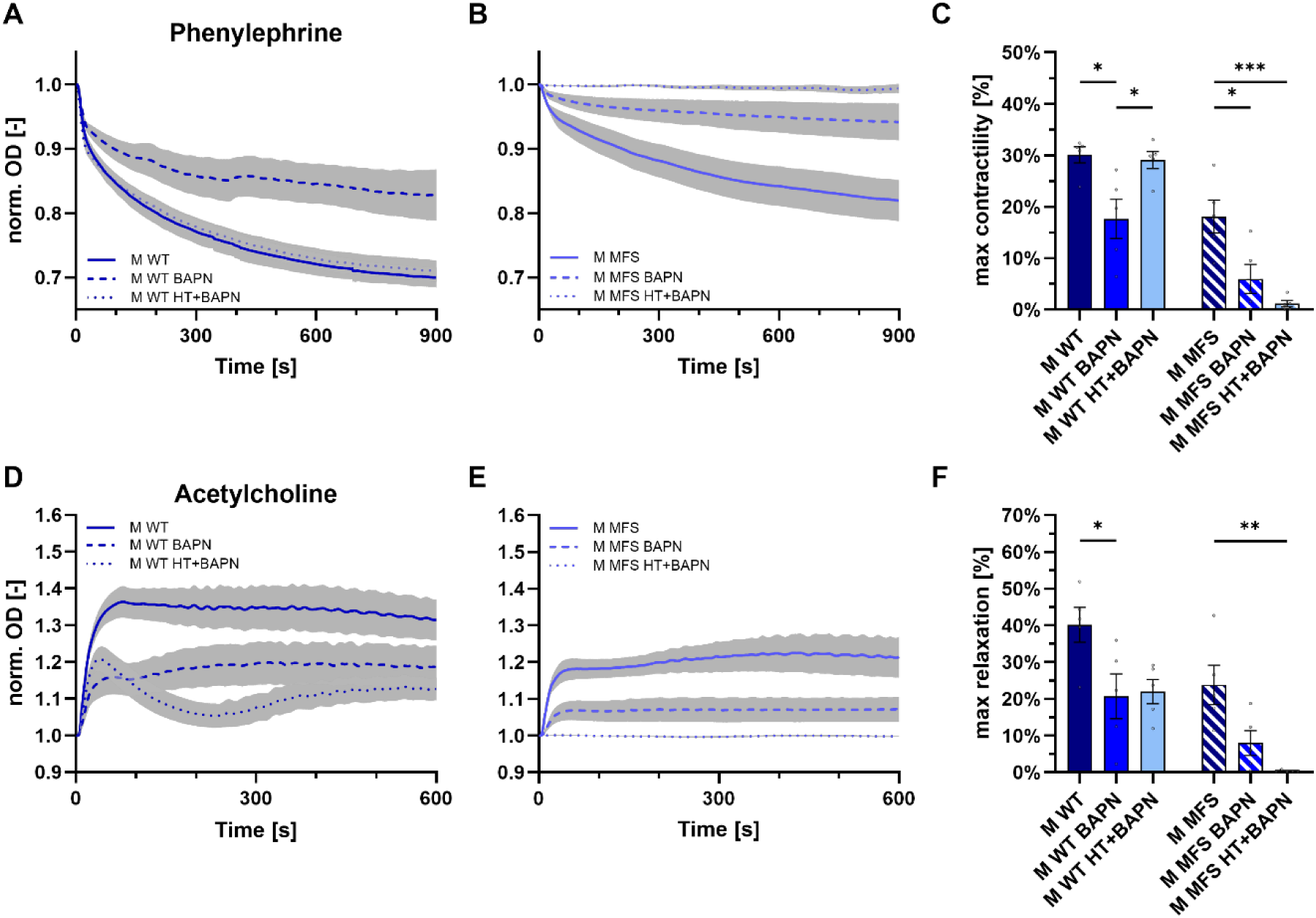
In male mice, BAPN reduces vasoactivity of both WT and MFS aortas while combined HT+BAPN renders male MFS mice inert. Vasoactive responses across six male study groups: normal, BAPN, and combined HT+BAPN. Panels A and B show time-course reductions in outer diameter in response to the vasoconstrictor phenylephrine for *Fbn1^+/+^* (WT) and *Fbn1^C1041G/+^*(MFS) mice respectively. Panels D and E show similar time-courses, this time for relaxation in outer diameter in response to the vasodilator acetylcholine. Panels C and F show the near steady state maximum contractile and relaxation response in all six groups. Data expressed as mean ± SEM. Significance only shown within a genotype, indicated by * (*p* < 0.05), ** (*p* < 0.005), and *** (*p* < 0.0005). See Supplemental Figure S3 for additional comparisons of vasoactive response.

Female WT aortas showed qualitatively similar vasoactive responses to the various stimulants as the male WT mice, though with a notable exception (**Supplemental Figures S4-S5**). While aortas in the female WT HT+BAPN group showed comparable evidence of SMC function as the female WT control mice (PE from 25.8% to 24.2%, KCl from 18.5% to 13.8%) and evidence of endothelial dysfunction (Ach from 29.9% to 10.5%, L-NAME from 23.1% to 7.2%), the vasoactive response of aortas in the female WT BAPN group was markedly greater than that of control mice across all agonists. The mechanism for this hyper-reactive phenomenon is unclear. This augmented vasoactivity in BAPN alone was preserved in the female MFS mice in response to PE and Ach, with nearly completely preserved contractile response to KCl and L-NAME. Notably, however, the female MFS mice given HT+BAPN experienced near-complete EC dysfunction (quantified by Ach and L-NAME response) though only minimally diminished SMC function (quantified by PE and KCl). Thus, male Marfan aortas were most susceptible to losses in vasoactive function while female WT and MFS aortas were partially resilient to both BAPN and HT+BAPN challenges.

### Passive Behavior

Material parameters for the four-fiber family constitutive relation were determined separately for each specimen, with values listed by group average (**Supplemental Table S2**) and individual specimen (**Supplemental Data File S1**). These parameters were used to compute multiple pressure-dependent mechanical metrics at both the group-specific systolic pressures (**Supplemental Tables S3-4**) and a common fixed pressure of 100 mmHg (**Supplemental Tables S5-6**). The male Marfan aorta experienced the most adverse remodeling in the BAPN groups, both without and with the added hypertensive challenge, as inferred by their dramatic changes in mechanical properties (**Figure 3**). By contrast, aortas from male WT mice nearly preserved their geometry and mechanical function in both the BAPN and HT+BAPN groups. Aortas from the female mice tended to follow similar trends, though with less disease severity (**Supplemental Figure S6**).

**Figure 3.**
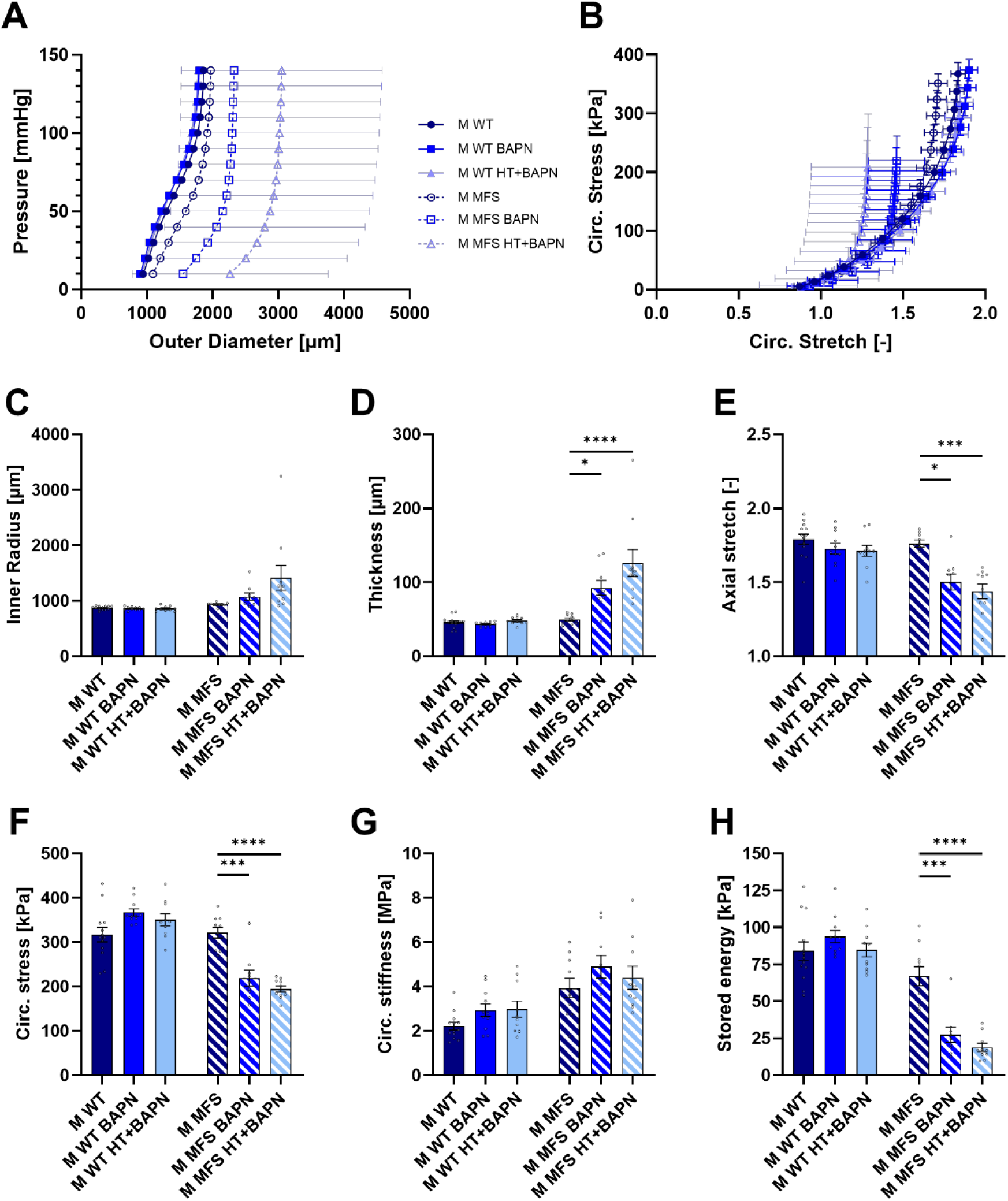
Male MFS mice showed the largest mechanical remodeling of the ascending aorta, resulting in marked changes in six key mechanical metrics. Both observed and computed mechanical metrics for six male study groups: no-diet, BAPN, and combined HT+BAPN; *Fbn1^+/+^* (WT) and *Fbn1^C1041G/+^* (MFS). Panels A and B show overall pressure-diameter response and the computed circumferential stress-stretch relationship. Panels C-E show distinct geometric remodeling while panels F-H show mechanical metrics computed from the four-fiber family constitutive equation. Data expressed as mean ± SEM. Significance only shown within a genotype, indicated by * (*p* < 0.05), *** (*p* < 0.0005), and **** (*p* < 0.0001). See Supplemental Figures S6-S8 for additional comparisons of passive mechanics.

Male WT aortas experienced only slight inward remodeling (<2% leftward shift in P-D relationship) in both the BAPN and HT+BAPN groups, namely minimal change in luminal inner radius (∼0% for BAPN, ∼2% decrease for HT+BAPN) with modest changes in wall thickness (∼7% decrease for BAPN, ∼4% increase for HT+BAPN). Meanwhile, male Marfan aortas experienced dramatic luminal dilatation (∼15% for BAPN, ∼51% for HT+BAPN) and significant wall thickening (84% for BAPN, ∼152% for HT+BAPN) as seen in **Figure 3a,c,d**. Male Marfan aortas also experienced a significant drop in the energetically preferred *in vivo* axial stretch (∼15% for BAPN, ∼18% for HT+BAPN) while reductions were less in male WT mice (∼2% for BAPN, ∼4% for HT+BAPN) (**Figure 3e**). The changes in male Marfan aortas suggested remodeling consistent with compromised mechanical homeostasis, which often results from altered mechanosensing or mechanoregulation of matrix, or both. A typically highly mechano-regulated metric is mean circumferential wall stress (**Figure 3f**), which depends on distending pressure, luminal radius, and wall thickness (Humphrey and Schwartz 2021; Matsumoto and Hayashi 1994). Male WT aortas experienced modest increases in circumferential stress (∼19% for BAPN, ∼10% for HT+BAPN), suggesting a degree of adaptive remodeling, while Marfan aortas experienced a marked drop in circumferential stress (∼32% for BAPN, ∼40% for HT+BAPN), suggesting maladaptive or over-exuberant remodeling. Ultimately, this remodeling of the male Marfan aorta resulted in an increase in circumferential stiffness (∼24% for BAPN, ∼12% for HT+BAPN) and marked loss of elastic energy storage capability (∼60% for BAPN, ∼72% for HT+BAPN), indicative of a loss of elastic function in the media (**Figure 3g-h**). Effects of circumferential stiffening and marked wall thickening resulted in a shift in the circumferential stress-stretch relationship, particularly in male MFS mice (**Figure 3b**). By contrast, male WT aortas experienced slightly higher circumferential stiffening (∼33% for BAPN, ∼35% for HT+BAPN) while maintaining stored energy within ∼15% of the NT control group.

Changes were similar in female aortas, albeit again to a lesser degree (**Supplemental Figure S6**). Interestingly, female Marfan aortas experienced distinctive thickening of the wall (∼27% for BAPN, ∼104% for HT+BAPN) and reductions in the *in vivo* axial stretch (∼5% for BAPN, ∼16% for HT+BAPN) in response to both BAPN and HT+BAPN but less luminal dilatation (∼9% for BAPN, ∼18% for HT+BAPN). This resulted in attenuated circumferential stiffening (∼24% for BAPN, ∼9% for HT+BAPN) and stored energy loss (∼19% for BAPN, ∼43% for HT+BAPN) compared to male counterparts. Similar to male WT aortas, female WT aortas experienced less wall thickening (∼17% for BAPN, ∼25% for HT+BAPN) and minimal luminal dilatation (∼4% decrease for BAPN, ∼2% increase for HT+BAPN). A variable response in circumferential stiffness (∼22% decrease for BAPN, ∼39% increase for HT+BAPN) was accompanied by a moderate drop in stored energy (∼18% for BAPN, ∼17% for HT+BAPN) in female WT aortas.

Additional comparisons across 10 key geometric and mechanical metrics are shown in Supplemental Materials for all WT (**Supplemental Figure S7**) and MFS (**Supplemental Figure S8**) groups, male and female. In summary, overall, Marfan aortas experienced more dramatic remodeling responses in geometry (luminal radius, thickness, *in vivo* biaxial stretch) and mechanical properties (biaxial stress and material stiffness as well as stored energy and distensibility) compared to WT mice, with increased severity in male compared with female MFS mice.

### Histology

Changes in function (vasoactivity and mechanical function) are generally accompanied by changes in form, including microstructural remodeling (**Figure 4, Supplemental Figures S9-S11, Supplemental Tables S7-S10**). Male MFS mice experienced the most dramatic changes in histological features, immediately evident by changes in medial vs. adventitia wall area extracted from MOV and PSR imaging. In male mice, BAPN was sufficient to induce a stepwise increase in medial wall area (WT ∼21%; MFS ∼34%) with a further increase with hypertensive challenge (WT ∼37%; MFS ∼39%). Adventitial wall area increased dramatically in male MFS mice (by ∼181% for BAPN; by ∼303% for HT+BAPN) while increasing only slightly in WT mice (by ∼13% for BAPN; by ∼19% for HT+BAPN). Within the male MFS groups, Spearman’s Rank correlation analysis revealed emerging correlations between adventitial area fraction and mechanical metrics, including circumferential stiffness (*r*=0.75, *p*=0.005), elastic stored energy (*r*=-0.77, *p*=0.002), distensibility (*r*=-0.86, *p*=0.000), and pulse-wave velocity (*r*=0.86, *p*=0.000). These findings would seem to be intuitive, as bolstering the adventitia through the deposition of collagens would typically stiffen the vessel and therefore reduce both distensibility and elastic function. The quality of newly deposited collagen in the presence of BAPN is unclear, however. Finally, male WT groups – which experienced minimal remodeling – showed no such correlations.

**Figure 4.**
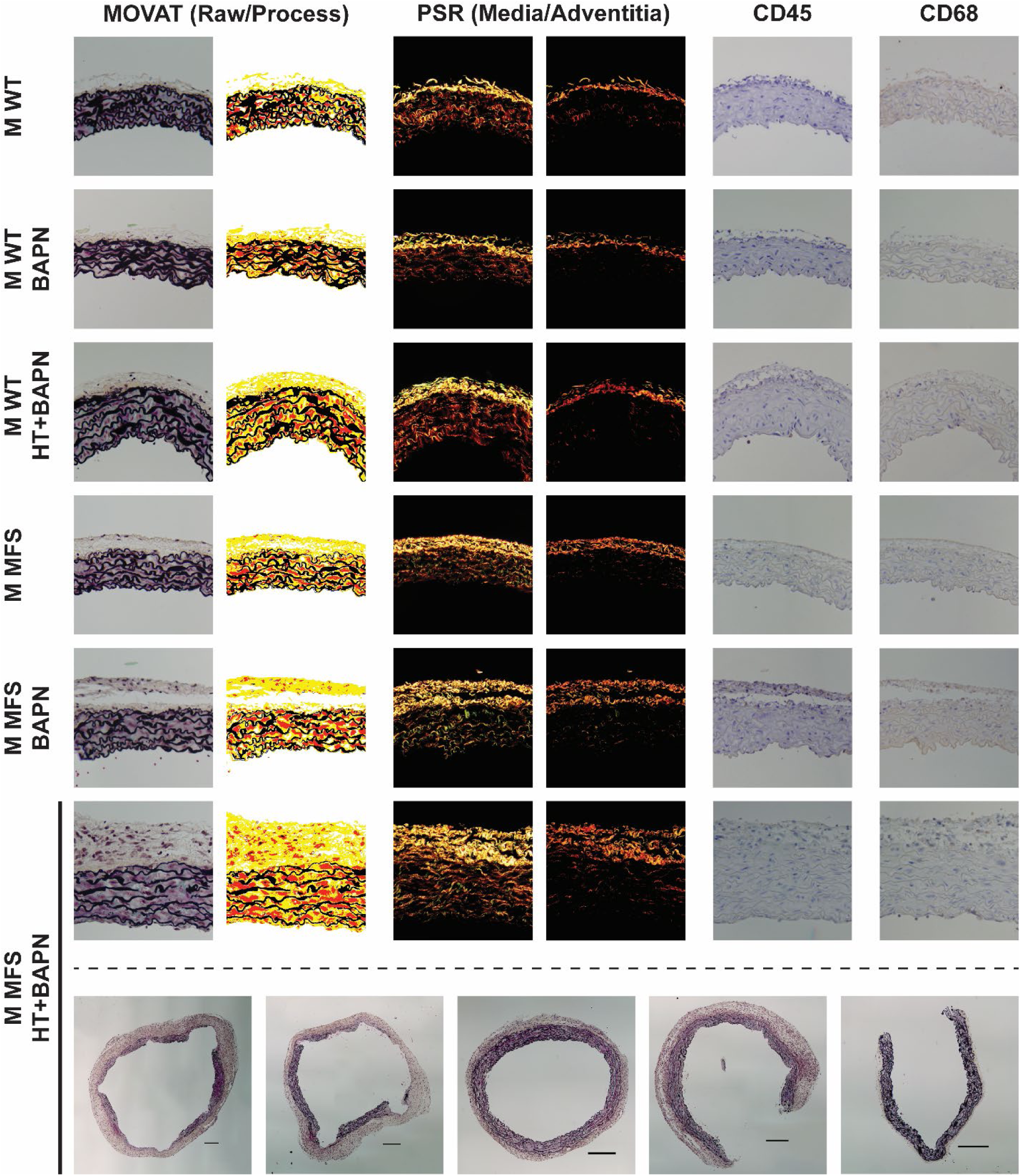
Representative histological analysis of male groups. Representative analysis of unloaded cross-sections including Movat’s pentachrome (pre- and post-processed sections), picrosirius red (captured at separate exposures to image specifically the media and adventitia), CD45, and CD68 staining. Note that the male MFS HT+BAPN groups is characterized by dramatic adventitial thickening and elastin fragmentation, with all five quantified samples shown in the bottom row. Representative sectioned images are shown in a 250 µm x 250 µm box. Scale bars on complete sections of male MFS HT+BAPN samples (bottom row) indicate a 200 µm length. See Supplemental Figure S9 for quantifications.

Immunohistochemistry revealed changes in CD45+ and CD68+ cells in male MFS mice but not WT mice. Interestingly, the highest change in positive expression occurred in male MFS mice with BAPN (CD45 from 0.05% to 0.54%; CD68 from 0.06% to 0.18%), rather than HT+BAPN (CD45 from 0.05% to 0.25%; CD68 from 0.06% to 0.10%). This could be a consequence of true reduced immune cell response in male MFS HT+BAPN or survival bias. Conversely, female MFS mice demonstrated no change in CD68 area fraction but an opposite trend to male MFS, in which CD45-positive area was reduced by BAPN (from 0.19% to 0.07%) but increased in HT+BAPN (from 0.19% to 0.48%). Nevertheless, CD45 and CD68 expressions were relatively low, and we thus focused on matrix constituents revealed by Movat’s pentachrome stain and picrosirius red: elastin, cytoplasm, and collagen in the media and mainly cytoplasm and collagen in the adventitia, noting that glycosaminoglycans (GAGs) and fibrin constituted a small portion of the wall area in all groups.

Medial elastin area varied by genotype. Male WT aortas experienced an increase in medial elastin area fraction for BAPN (from 44.2% to 55.3%) and HT+BAPN (from 44.2% to 49.0%), both with an associated near-identical increase in medial elastin area (∼0.52-fold increase), suggesting that addition of HT negligibly affected elastin area. By contrast, BAPN alone induced an expansion (likely de-compaction) of elastic lamellae. Male MFS mice experienced a differential response: BAPN increased medial elastin area (∼0.17-fold increase) while HT+BAPN decreased elastin area (∼0.20-fold decrease). Qualitative examination revealed in the male MFS HT+BAPN group that loss of elastin area associated with marked elastic fiber fragmentation, especially in the cases of dissection (bottom row of **Figure 4**). In both male genotypes, correlations emerged between mechanical metrics and medial elastin area fraction but not medial elastin area, suggesting that mechanical changes were not dictated by changes to medial elastin, but rather the observed increases in cytoplasm and collagen areas. Medial and adventitial cytoplasm area and area fraction increased in both male genotypes, albeit to a lesser degree in WT mice. Male WT mice experienced moderate increases in medial (∼0.15-fold for BAPN; ∼0.20-fold for HT+BAPN) and adventitial (∼0.70-fold for BAPN; ∼0.43-fold for HT+BAPN) cytoplasm area with minimal correlation with vasoactive and mechanical properties. Representative of dramatic remodeling, Male MFS experienced greater increases in medial (∼0.50-fold for BAPN; ∼1.15-fold for HT+BAPN) and adventitial (∼6.41-fold for BAPN; ∼12.64-fold for HT+BAPN) cytoplasm area. Here, moderate negative correlations emerged between medial cytoplasm area and vasoactivity (KCl *r*=-0.42, *p*=0.127; PE: *r*=-0.57, *p*=0.037; Ach: *r*=-0.57, *p*=0.037; L-NAME: *r*=-0.68, *p*=0.009), suggesting that increased cellular material need not contribute directly to vasoactivity, but instead may reflect an increased synthetic phenotype. This postulate is supported by the marked deposition of collagens in the male MFS groups.

### Pressure-Burst

Recall that several groups – particularly male MFS mice challenged with HT+BAPN – experienced premature deaths, some due to thoracic aortic rupture. Among the surviving mice, *in vivo* dissections and spontaneous *ex vivo* delaminations were scattered across male and female groups having some form of BAPN, with prevalence of *in vivo* dissections in the male MFS mice with HT+BAPN. Both contained (included dissection or delamination) and transmural (leading to death) failures occur when wall stress exceeds wall strength, the latter of which may be estimated using pressure-burst testing. Generally, BAPN and HT+BAPN reduced failure strength (i.e., lower circumferential stress at rupture) in both male and female mice when compared to control groups (**Figure 5**, **Supplemental Figure S12, Supplemental Table S11**). Male WT aortas thus ruptured at lower values of burst pressure for BAPN (by ∼25%) and HT+BAPN (by ∼27%), accompanied by minimal changes in burst circumferential stretch (∼1% for BAPN, ∼5% for HT+BAPN) (**Figure 5a-b**). Computation of circumferential stress at failure reveals a stepwise drop for BAPN alone (by ∼21%) then HT+BAPN (by ∼37%), likely influenced by the small degree of wall thickening and reduced dilatation of the HT+BAPN group (**Figure 5c**). By contrast, BAPN alone was sufficient to cause marked decreases in failure strength in male Marfan aortas, with variable drops in burst pressure (∼15% for BAPN, ∼9% for HT+BAPN) and circumferential stretch (∼7% for BAPN, ∼21% for HT+BAPN), ultimately resulting in a comparable reduction in circumferential stress at failure (∼43% for BAPN, ∼46% for HT+BAPN). That is, all male MFS mice exhibited lower circumferential stress at failure in comparison to the corresponding male WT mice, indicating a detrimental effect of the *Fbn1* variant on failure strength.

**Figure 5.**
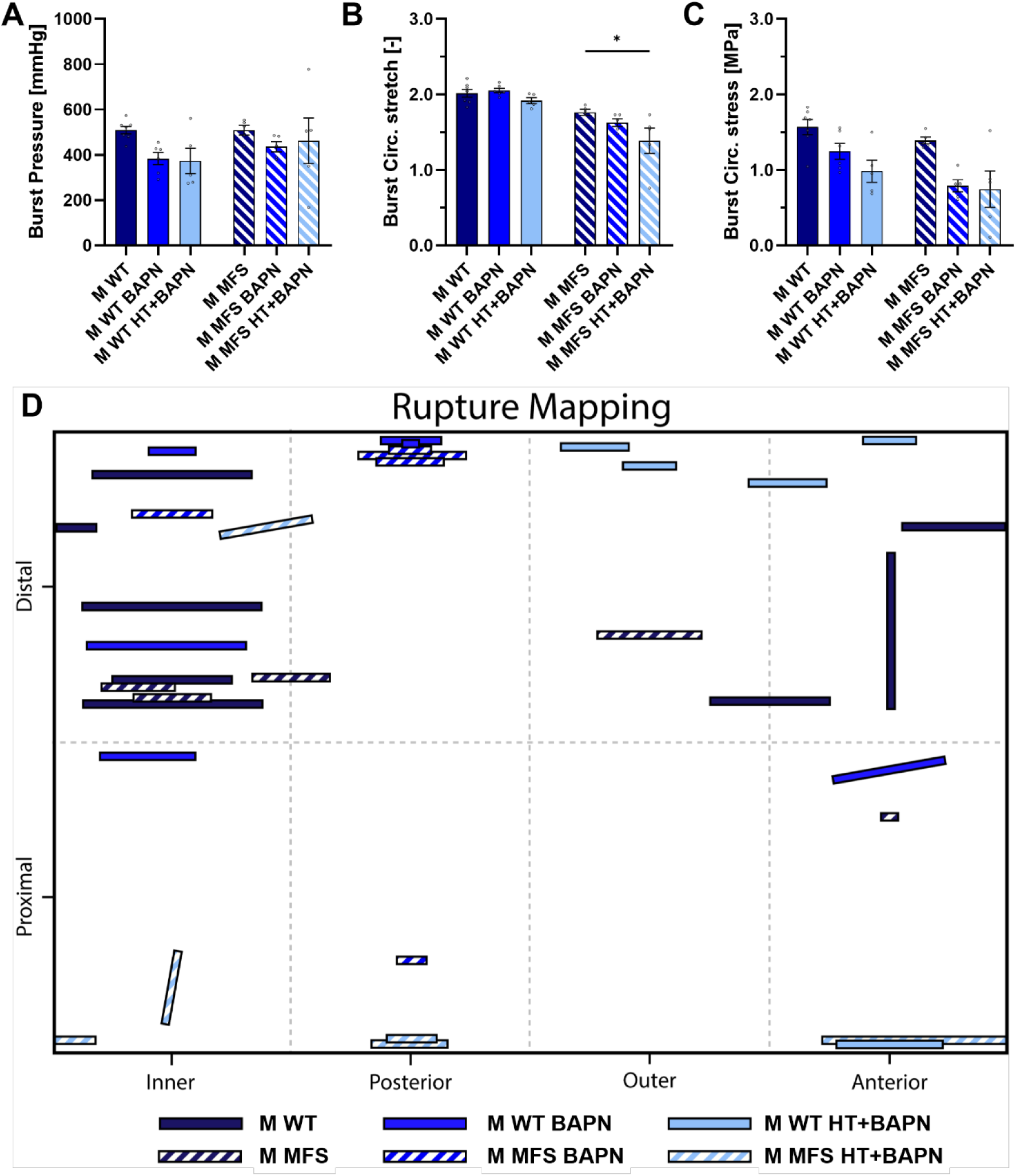
BAPN and HT+BAPN did not induce significant changes in rupture properties; however, a shift in location of rupture indicates potential localized vulnerabilities. Rupture mechanical analysis of six male study groups: no-diet, BAPN, and combined HT+BAPN; *Fbn1^+/+^* (WT) and *Fbn1^C1041G/+^* (MFS). Few significant changes occur in measured burst pressure (A), burst circumferential stretch (B), and computed circumferential stress at rupture (C). Panel D shows qualitatively the location, size, and orientation of rupture, noting a shift from distal regions to the aortic root in male MFS HT+BAPN mice. Data expressed as mean ± SEM. Significance only shown within a genotype.

While none of the measured or computed failure metrics were significantly different within each genotype, qualitative mapping of rupture sites showed a preference for the middle of the ascending aorta in control NT groups that differed by genotype (**Figure 5d**). Generally, male WT aortas demonstrated a distally moving rupture site from BAPN to HT+BAPN groups, eventually favoring the outer curvature at the base of the brachiocephalic artery —a site of pre-stress during vessel alignment for biaxial testing. Meanwhile, male Marfan aortas demonstrated a variable response for BAPN and HT+BAPN; BAPN alone led to rupture at the base of the brachiocephalic (similar to male WT mice) whereas HT+BAPN induced a preference for failure at the aortic root, which often coincided with the general site of *in vivo* dissection.

In contrast to findings in male mice, BAPN alone was sufficient to reduce failure strength in both female WT and MFS mice (**Supplemental Figure S12, Supplemental Table S11**). In female WT aortas, a comparable reduction in burst pressure (∼24% for BAPN, ∼23% for HT+BAPN) and burst outer diameter (∼6% for BAPN, ∼7% for HT+BAPN) resulted in a similar reduction in failure stress (∼39% for BAPN, ∼42% for HT+BAPN). Female MFS aortas also experienced a BAPN-induced reduction in failure properties; however, the wall thickening induced by the added hypertensive challenge ameliorated some degree of change in circumferential stress at burst. That is, while BAPN reduced the burst pressure in female Marfan aortas (by ∼27%) and increased the burst outer diameter (∼5%), the added hypertensive challenge increased both the burst pressure (∼5%) and burst outer diameter (∼18%). Together, these resulted in a ∼34% reduction in circumferential stress at failure for BAPN vs. a ∼29% reduction for HT+BAPN. Generally, all female groups demonstrated a higher factor-of-safety (failure stress / systolic stress) compared to the male groups, with the exception of the MFS BAPN group, which may reflect the higher 12-week survival rate in females. In contrast to males, all female groups tended to exhibit a similar rupture site near the distal portion of the inner curvature or anterior face. This site coincides with pre-stress associated with straightening of the ascending aorta curvature prior to biaxial testing.

### Necropsy Analysis

Noting again that biomechanical testing and analysis was necessarily restricted to mice that survived to the 12-week endpoint, resulting in an inherent survivorship bias that particularly impacted evaluation of the male MFS HT+BAPN group. Hence, it is best to consider this particular group as “aortas that remodeled sufficiently in response to HT+BAPN challenge”. To begin to elucidate driving factors of premature death in “aortas that remodeled insufficiently”, we performed histological analysis of all necropsied male MFS HT+BAPN samples that exhibited thoracic aortopathy at the time of death (**Supplemental Figure S13**). Qualitative examination of necropsies within the first seven days revealed frequent blood and/or fibrin deposits within the vessel wall. Following the first week of HT+BAPN, blood deposits largely disappear, instead giving way to evidence of elastin fragmentation and often large medial discontinuities. These may indicate evidence of post-wound healing in which intramural blood has been resorbed and discontinuities in elastin have been bolstered with newly deposited collagen, compromising biomechanical function while attempting to retain structural strength. These adaptive remodeling attempts ultimately failed, however, resulting in premature mortality.

## DISCUSSION

Under normal conditions, ECM synthesis is highest within the thoracic aorta during the early postnatal period, peaking in WT mice at about postnatal day P10 for elastin and P21 for collagen (Weiss et al. 2024). For this reason, administration of BAPN for short periods (2-4 weeks) typically has a much more dramatic effect on the aorta in younger than older mice and rats (Zheng et al. 2020; Sawada et al. 2022). Illustrative studies that have exploited the postnatal developmental period to drive BAPN-induced aortopathy in mice often started BAPN at 3- (Zhou et al. 2025) or 4- (Dar Weiss et al. 2023) weeks of age. Similar studies in juvenile rats reveal many important findings, including an overall increased aortic dilatation with decreased wall strength, decreased connections between SMCs and elastic fibers, and an increased accumulation of GAGs (Hosoda and Iri 1966; Brüel et al. 1998). Herein, we gave BAPN (2.0 g/L in drinking water) to adult normotensive male and female WT and MFS mice from 8 to 12 weeks of age both to verify additional results from a prior study (Weiss et al. 2023) and to facilitate direct comparison with our planned HT+BAPN groups. The two primary differences between the former and present BAPN-only study were the less frequent refreshing of the drinking water in the prior study (1x per week vs. 3x herein) and the added evaluation of vasoactivity herein. Results were qualitatively similar between studies, but changes were quantitatively greater herein. When evaluating passive biomechanical metrics for the male Marfan aorta at a common pressure of 100 mmHg, in the present study BAPN caused greater increases in luminal radius (16.6% vs. 9.1% previously), wall thickness (83.3% vs. 41.4%), and circumferential stiffness (23.7% vs. 9.5%) as well as greater decreases in energetically preferred axial stretch (14.7% vs. 5.1%), circumferential wall stress (33.3% vs. 14.4%), and elastic energy storage (58.0% vs. 21.6%). These findings implicate a temporal decrease in BAPN potency within drinking water, and thus a need to detail the methods used (cf. Daugherty et al. 2025). The present BAPN-only study further revealed qualitatively similar yet quantitatively more severe changes in biomechanical characteristics than in our prior hypertension-only study in MFS mice (Mays and Humphrey 2026), which elevated blood pressure using the same high salt diet (8% NaCl) and L-NAME (3.0 g/L in drinking water replenished 3x weekly). At a common evaluation pressure of 100 mmHg, the prior hypertension-alone study in male MFS mice demonstrated more modest increases in luminal radius (9.4%) and wall thickness (70.0%) accompanied by decreases in energetically preferred axial stretch (14.7%), circumferential wall stress (29.7%), and elastic energy storage (41.6%).

Although BAPN has been combined with hypertension during postnatal development to great effect (Nakashima and Sueishi 1992), we reasoned that effects of LOX inhibition would be similarly dramatic in hypertensive adult mice since initial increases in wall stress resulting from increasing blood pressure decrease collagen half-life (Nissen et al. 1978) and increase rates of ECM synthesis (Spronck et al. 2021). Indeed, a prior study co-delivered BAPN during the first 2 weeks of a 6-week chronic angiotensin II infusion study in initially 9-week-old male C57BL/6J mice (Kanematsu et al. 2010). The investigators reported a 71% incidence of ruptured or unruptured aortic aneurysm and 2% incidence of dissecting aneurysm. Conversely, none of the mice given either BAPN or angiotensin II alone developed an aneurysm, suggesting that blocking cross-linking of newly synthesized ECM during periods of increased ECM turnover (i.e., combined hits) work together to increase aortopathy following vascular maturity. Surprisingly, biomechanical characterization remained absent when combining BAPN and hypertension, with prior studies favoring morphological and histological examination. Biomechanics can be especially insightful when examining severe aortic phenotypes since it is axiomatic that failure (i.e., dissection and/or rupture) occurs when wall stress exceeds wall strength.

Most importantly herein, combining hypertension and BAPN from 8 to 12 weeks of age resulted in severe thoracic aortopathy in male *Fbn1^C1041G/+^* mice, characterized by a > 50% dilatation relative to age-, sex-, and genotype-matched normotensive MFS controls (and > 60% relative to WT controls), a 70% incidence of *in vivo* dissection (vs. 10% for similarly challenged male WT mice), and an ∼67% mortality (compared to 22% for similarly challenged male WT mice). There were, in addition, significant changes in active and passive biomechanical properties, reflective of marked decreases in aortic functionality, that were consistent with altered histological metrics. In brief, the induced hypertension (high salt + L-NAME) increased the baseline mechanical load on the media (largely elastin, but also collagen and SMCs) and adventitia (mainly collagen), prompting an increased turnover of ECM. Concurrent BAPN delivery blocked the compensatory upregulation of LOX in MFS (Busnadiego et al. 2015), causing an apparent catastrophic reduction of ECM cross-linking. Thus, the adventitia thickened with immature (i.e. insufficiently crosslinked) collagen, stiffening the aortic wall structurally (increased PWV) while driving down circumferential stress (a key mechano-stimulus for synthesis). Moreover, there was a near-complete loss of vasoactive function (**Figures 2, S3**) as well as elastic stored energy (**Figure 3**), likely caused in large part by the medial elastin fragmentation found under histological examination (**Figure 4**). Fragmentation of elastic lamellae and interlamellar “struts” which SMCs adhere to and actuate upon drive SMC anoïkis and subsequent loss of vasoactive capabilities of the artery (Michel 2003). The increased cellular material in the media of these critically fragmented vessels would likely adopt a synthetic phenotype, as adaptive remodeling attempts continue. Critically, there was also a marked reduction in circumferential strength. This matrix remodeling in the male MFS HT+BAPN mice not only changed bulk material properties, it appeared to induce local structural vulnerabilities that resulted in increased rates of *in vivo* dissections or *ex vivo* delaminations (**Figure 1**), particularly about the aortic root (**Figure 5**).

From our biomechanical analysis we can infer that the 12-week-old MFS mice following HT+BAPN challenge remain at high risk of rupture, and extension of diets beyond 28 days will result in continued mortality. While the previously evaluated hypertensive challenge (Mays and Humphrey 2026) induced high mortality of male MFS within the first 14 days of diet in tandem with step-wise increases in blood pressure, the present combined hypertension and BAPN challenge induces mortality rates of ∼3.6% mortality/day within the first 0-14 days followed by ∼1.5% mortality/day in the latter 15-28 days and no step-wise increase in blood pressure (**Figure 1**). The lack of apparent blood pressure increases exceeding 140 mmHg – as previously seen in hypertensive challenge alone – may be due to (i) adaptive response of surviving mice, (ii) elimination by premature death of all mice that experienced acute blood pressure elevation, or (iii) interactive response between the hypertension and BAPN. Ultimately, the end-point HT+BAPN phenotype displayed biomechanical characteristics of late-stage aging, exceeding those of 2-year-old *Fbn1^C1041G/+^* mice nearing end-of-life (Means et al. 2026), including greatly diminished vasoactivity (HT+BAPN: PE 1.1%, Ach 0.3%, 2-year-old: PE ∼5.6%, Ach ∼4.8%), circumferential stiffening (HT+BAPN: 4.40 MPa, 2-year-old: 3.13 MPa), and loss of stored energy capability (HT+BAPN: 19 kPa, 2-year-old: 50 kPa) at systolic pressures.

The role of even mild pre-existing *Fbn1* dysfunction in MFS mice in predisposing to severe aortopathy cannot be overstated, as littermate wild-type (*Fbn1^+/+^*) mice remained profoundly resilient against HT+BAPN challenge, often expressing a less severe phenotype than for HT alone (cf. Mays and Humphrey 2026) while maintaining high survival rates. When compared to non-treated normotensive (NT) controls, the HT, BAPN, and HT+BAPN induced only modest changes in WT mice in luminal radius (−8% to HT; −2% to BAPN; −6% to HT+BAPN), wall thickness (+12% to HT; −4% to BAPN; +9% to HT+BAPN), circumferential stress (−18% to HT; −1% to BAPN; −15% to HT+BAPN), and circumferential stiffness (−22% to HT; −3% to BAPN; −16% to HT+BAPN). Computing luminal radius and wall thickness together reveals a ∼3% increase in loaded cross-sectional area to HT+BAPN – a near-negligible change, especially when compared against the ∼292% increase experienced by MFS HT+BAPN mice. Interestingly, HT alone resulted in the greatest loss of SMC (PE from 30.1% to 23.8% in HT vs. 24.3% in BAPN and 29.1% in HT+BAPN) and EC (Ach from 40.1% to 11.3% in HT vs. 35.2% in BAPN and 22.0% in HT+BAPN) function. While functional changes by ECs was expected in cases of chronic L-NAME delivery, near-complete preservation of SMC contractility in WT mice under combined HT+BAPN challenge was surprising. Nevertheless, despite the markedly reduced remodeling as compared to MFS mice, BAPN yet reduced the integrity of newly deposited matrix constituents in WT mice. This can be seen in PSR staining as differences in adventitial collagen area (+61% to HT vs. +38% to BAPN and +42% to HT+BAPN) and circumferential failure stress (+26% to HT vs. −21% to BAPN and −37% to HT+BAPN). The reduction of competent cross-linking of newly deposited matrix during adaptive remodeling is indeed what drives the compromised MFS HT+BAPN mouse towards such distinctive aortopathies.

Female mice exhibited increased survival and more adaptive biomechanical remodeling responses as compared to corresponding male groups. A notable exception is the F WT HT+BAPN group, in which female mice experienced greater *in vivo* dissection (17% vs 10% in males) and *ex vivo* delamination (25% vs 0%) rates, which may be attributed to the higher survival rate (92% vs 78%) allowing for the development of complex aortopathy. Consequently, at a fixed pressure of 100 mmHg, the F WT HT+BAPN group experienced a ∼43% increased circumferential stiffness (1.63 vs 1.14 MPa) and ∼67% increased adventitial collagen area (∼77,900 vs ∼46,700 µm^2^) though reduced vasoreactivity (PE: 24.2% vs 29.1%; Ach: 10.5% vs 22.0%) when compared to the M WT HT+BAPN group. Conversely, all female MFS groups maintained both a higher survival rate and greater evidence of mechanical homeostasis. In particular, the F MFS HT+BAPN group remodeled less (∼16% lower thickness, ∼30% lower lumen radius), stiffened less (∼41% lower circumferential stiffness, ∼39% lower adventitial collagen area), maintained a ∼68% higher stored energy (27 kPa vs 16 kPa), and retained some degree of vasoactive function (PE: 16.7% vs 1.1%; Ach: 6.5% vs 0.3%) compared to the M MFS HT+BAPN group. Such distinctive sex differences in MFS have been previously noted in mice (Gharraee et al. 2022; Dar Weiss et al. 2023; Means et al. 2026) and humans (Roman et al. 2017), though further mechanistic investigation is needed.

Notwithstanding lessons learned, this study is not without limitations. There are differential responses along the aorta to both hypertension and BAPN (Bersi et al. 2017; Franklin et al. 2024), but we focused on the ascending aorta since it is the most affected segment in MFS. Much could be learned from studies of other vascular segments. One should similarly study possible regional differences in the mechanics (Seshaiyer and Humphrey 2003; Bersi et al. 2016), especially given the geometric and loading heterogeneity of the ascending aorta. Marked differences in aortopathy arise depending on BAPN concentration (Franklin et al. 2024); we considered only one concentration of L-NAME and BAPN. Others have shown therapeutic benefit of different drugs (Busnadiego, Gorbenko del Blanco, et al. 2015; Kanematsu et al. 2010; B. Zhou et al. 2019), but we did not consider potential pharmacological interventions. There are advantages to collecting longitudinal data (e.g., at 1, 2, and 4 weeks) to reduce the impact of survivorship bias as well as to inform computational models of the growth and remodeling processes (Humphrey 2021), but we focused on one duration given the many different groups (12) and types of testing (biomechanics, histology, sequencing). Mice in the HT+BAPN group had reduced body mass. Allometric scaling (cf. Korneva et al. 2019) could further inform computational models. The pressure-burst testing was conducted without measuring axial force at rupture. Given that most ruptures aligned circumferentially, axial force measurement would have allowed computation of axial stress at failure. Finally, we observed a strong sexual dimorphism but did not attempt to identify underlying mechanisms (e.g., adding ovariectomized mice).

The *Fbn1^mgR/mgR^* mouse model of MFS is widely studied given its severe phenotype, with marked aortic dilatation and 50% mortality by 9 weeks of age in the absence of genetic drift. Yet, this mouse does not model a human pathogenic variant or many of the associated differentially expressed genes (Sun et al. 2023). By contrast, the *Fbn1^C1041G/+^* mouse models a heterozygous missense variant found in patients, but it typically exhibits such a mild phenotype that utility is otherwise limited. We showed that a severe phenotype can be induced in adult male *Fbn1^C1041G/+^* mice by combining hypertension with an inhibitor of lysyl oxidase. Whereas others have used infusion of angiotensin II to increase blood pressure in mice, this method can induce thoracic aortic disease independent of pathogenic variants (Rateri et al. 2014; Bersi et al. 2017b), thus we sought to use a more relevant method to increase blood pressure (high salt diet and endothelial dysfunction). Given that there is an increase in lysyl oxidase in the aorta of MFS mice (Busnadiego et al. 2015; Weiss et al., 2023), we further sought to block this compensatory response. Finally, since most disease progression in human MFS patients results in interventions in adults, often in their 30s or 40s (Jondeau et al. 2012), we sought to accelerate the phenotype in adult mice.

Motivated thus, the current findings confirmed sexual dimorphism in Marfan mice, with hypertension worsening the disease phenotype but potentially eliciting a compensatory deposition of collagen that could be protective if cross-linked properly. When cross-linking was blocked, even the marked reduction in wall stress resulting from a thickened wall did not compensate for the marked reduction in wall strength. Together, these data suggest that lowering wall stress with anti-hypertensive medications can provide a protective benefit but there is a need for robust matrix deposition and organization. Computational methods, such as growth and remodeling models (Humphrey 2021), could provide further insights into roles of differential mechanical loading and matrix remodeling in rendering the aorta more or less vulnerable to dissection and rupture. We submit that the current data will better inform such models. Similarly, experimental investigations into possible interventions that increase compensatory reparative processes within the aortic wall should be pursued while avoiding any medications that may compromise collagen integrity (e.g., ciprofloxacin; LeMaire et al. 2022, 2026).

## ACKNOWLEDGMENTS

This work was supported, in part, by grants from the US National Institutes of Health (R01 HL169147, P01 HL169168 to JDH) and the Leducq Foundation (erAADicate).

