## Supplemental Figures and Tables for "Blocking Compensatory Matrix Cross-Linking Accelerates Thoracic Aortopathy in a Mouse Model of Marfan Syndrome"

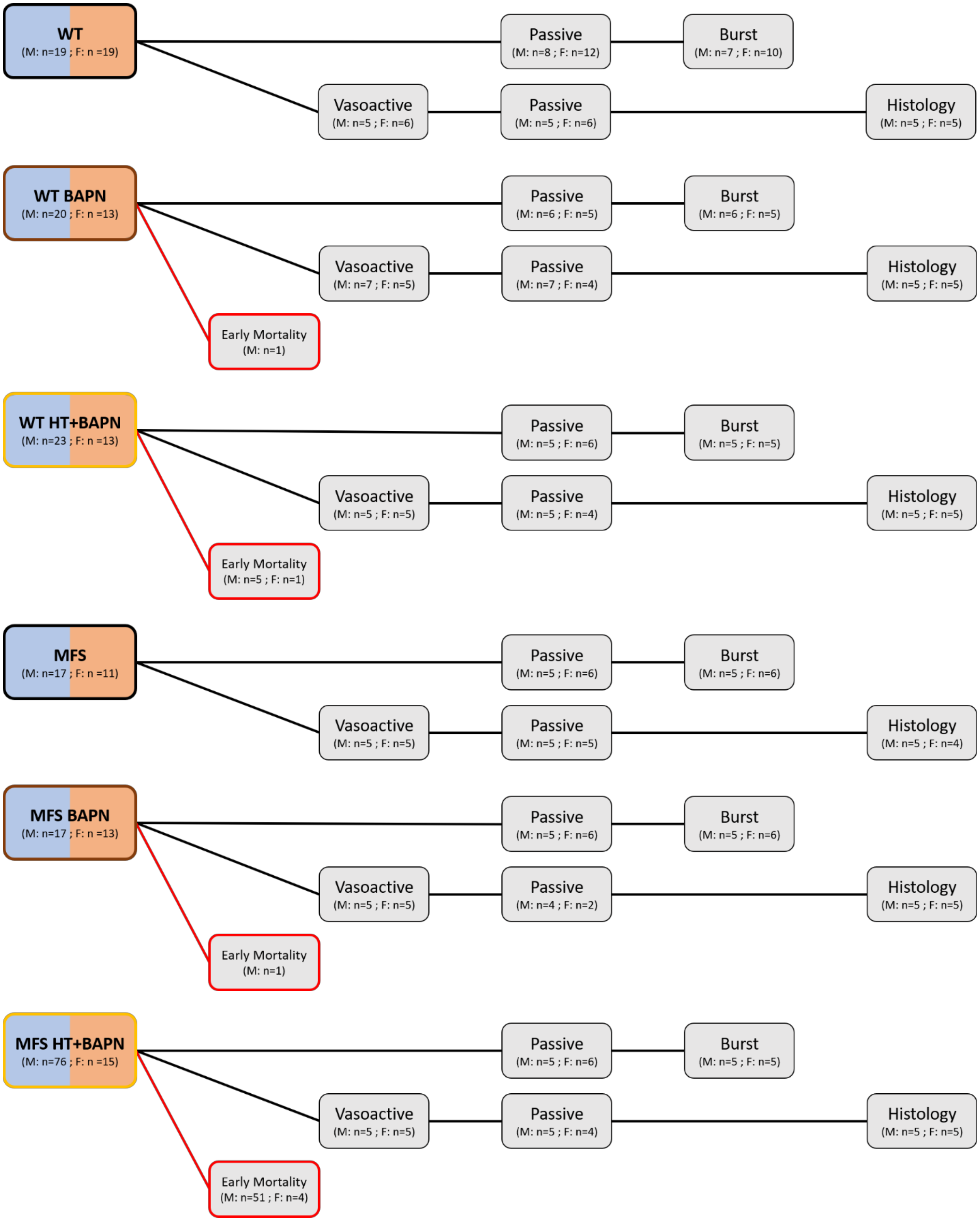


**Flowchart S1. Detailed accounting of allocation for all 256 mice, 171 of which were used for biaxial testing.** Shown are numbers of male (M) and female (F) littermate *Fbn1*$^{+/+}$ (WT) and *Fbn1*$^{C1041G/+}$ (MFS) mice allocated towards vasoactive, passive, and burst biaxial studies. Numbers in red boxes indicate mice removed from consideration for biaxial testing due to early mortality.

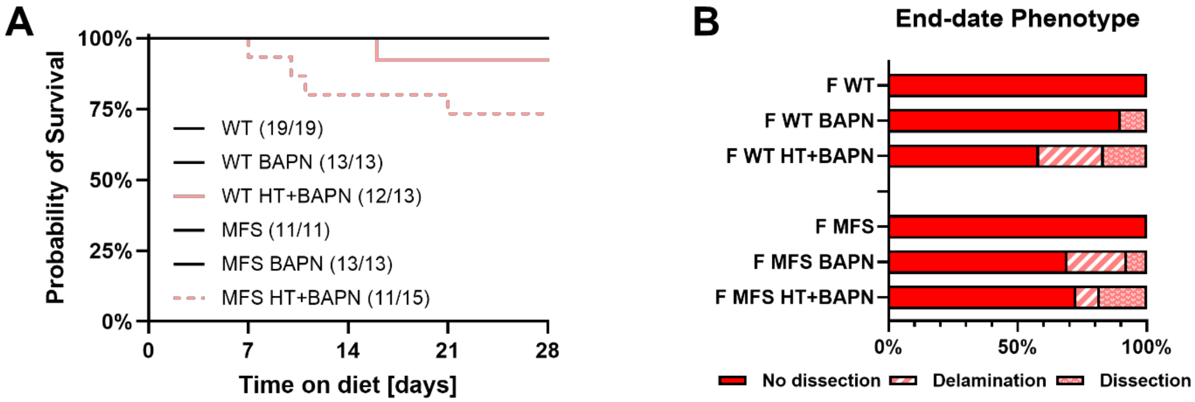


**Supplemental Figure S1. Female groups experienced milder phenotypes than male mice after four weeks of BAPN and HT+BAPN.** Combined HT+BAPN was necessary to induce early mortality in female mice (A). BAPN alone was sufficient to induce a dissection phenotype in female mice (B), although ultimately in lesser numbers than in male mice.

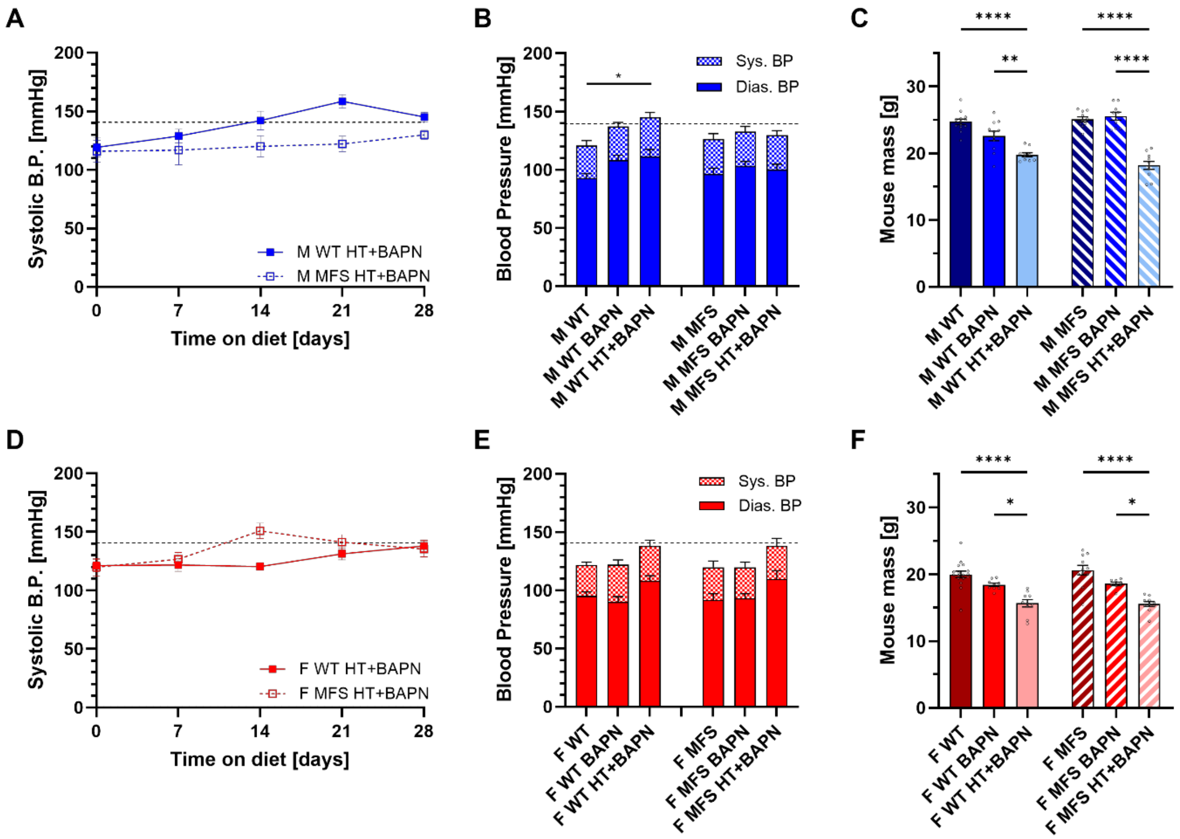


**Supplemental Figure S2. Blood pressure gradually elevated in both sexes and genotypes in response to HT+BAPN accompanied by significant losses in body mass**. Tail-cuff measured systolic blood pressure in male (M, blue) and female (F, red), *Fbn1*$^{+/+}$ (WT) and *Fbn1*$^{C1041G/+}$ (MFS) measured at weekly increments throughout the four-week study (A, D) and final values at 12-weeks of age (B, E). Combined HT+BAPN induced a significant loss of body mass in both sexes and genotypes (C, F). Data expressed as mean ± SEM. Significance only shown within a genotype, indicated by * ($p < 0.05$), ** ($p < 0.005$), and **** ($p < 0.0001$).

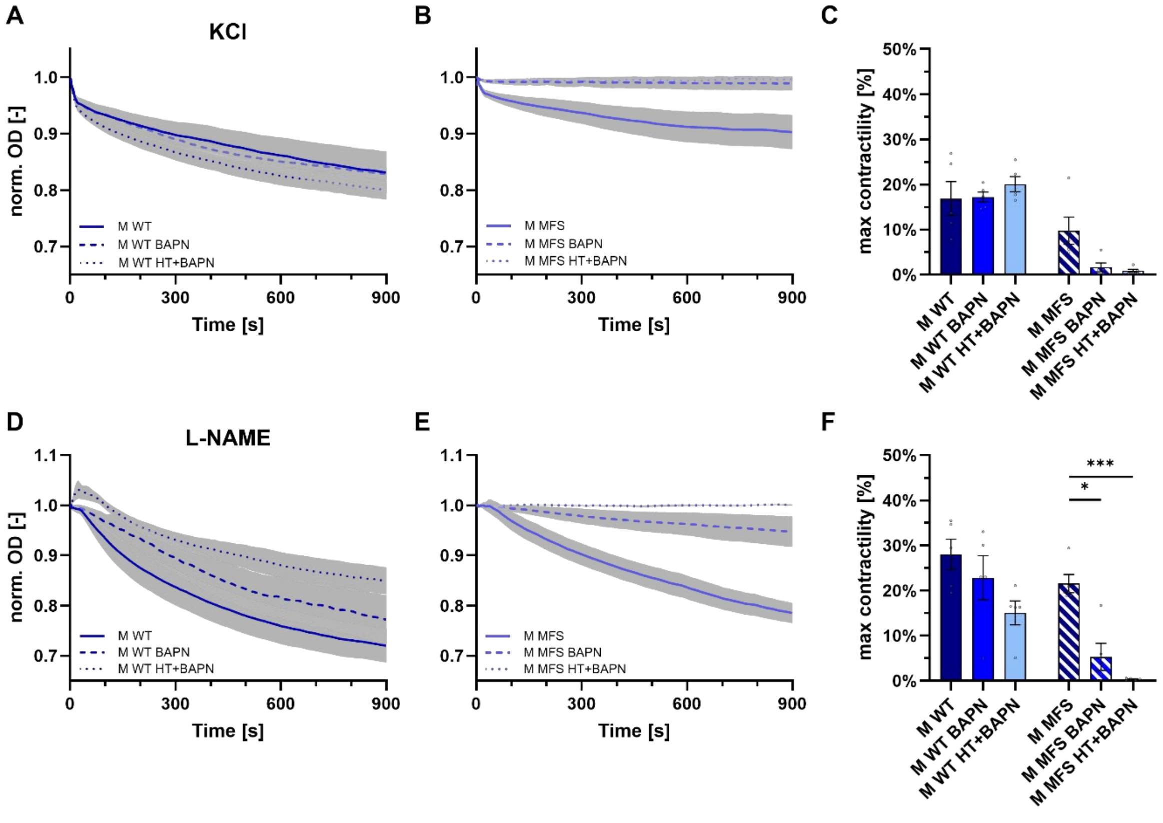


**Supplemental Figure S3. Vasoactive response to KCl and L-NAME showed similar loss in vasoactive function in male MFS mice.** Similar to Figure 2 in the main text except for vasoactive response to the vasoconstrictors potassium chloride (KCl) and L-NAME, an eNOS inhibitor. Panels A, B, D, and E show time-course reductions in outer diameter in response to agonists while panels C and F show near steady-state maximum changes in outer diameter. Data expressed as mean ± SEM. Significance only shown within a genotype, indicated by * ($p < 0.05$) and *** ($p < 0.0005$).

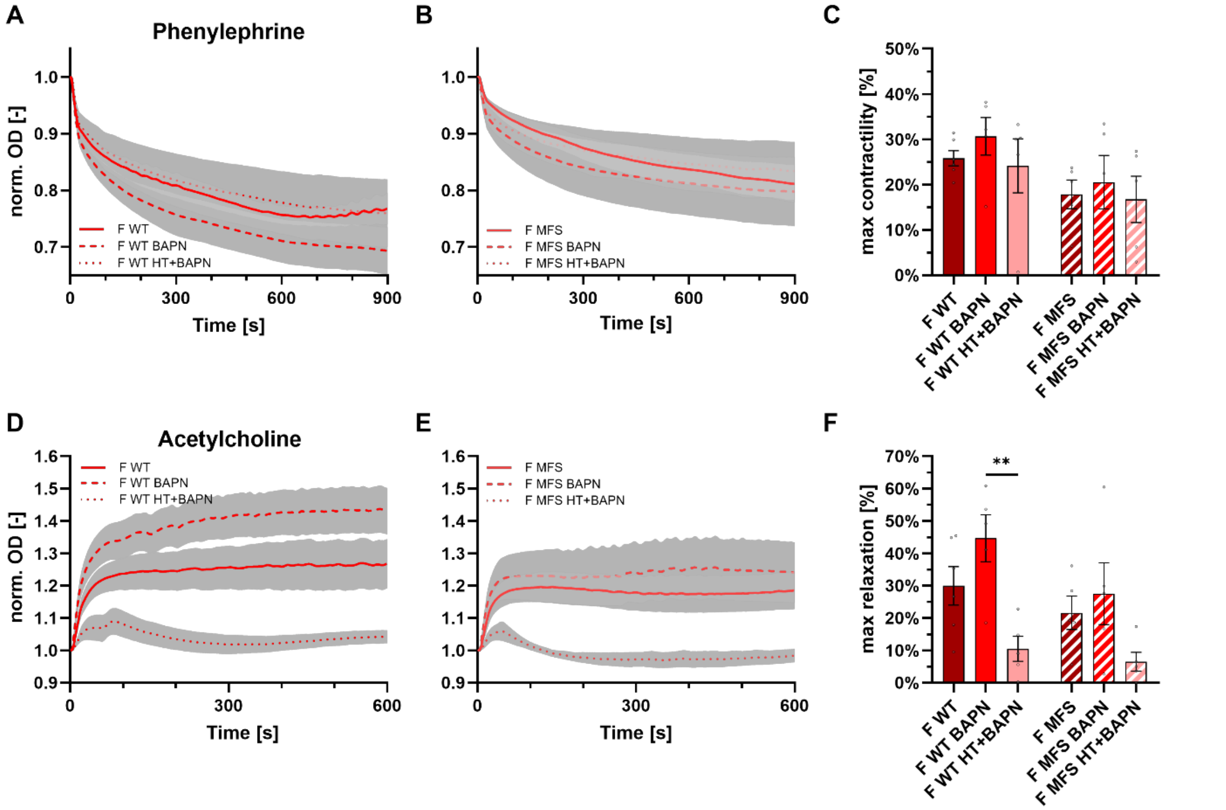


**Supplemental Figure S4. Vasoactive function of female mice was remarkably maintained with the exception of endothelial dysfunction in response to combined HT+BAPN challenge.** Similar to Figure 2 in the main text, except for vasoactive response to phenylephrine (panels A-C) and acetylcholine (panels D-F) in female mice. Panels A, B, D, and E show time-course changes in outer diameter in response to agonists while panels C and F show near steady-state maximum changes in outer diameter. Data expressed as mean $\pm$ SEM. Significance only shown within a genotype, indicated by ** ($p < 0.005$).

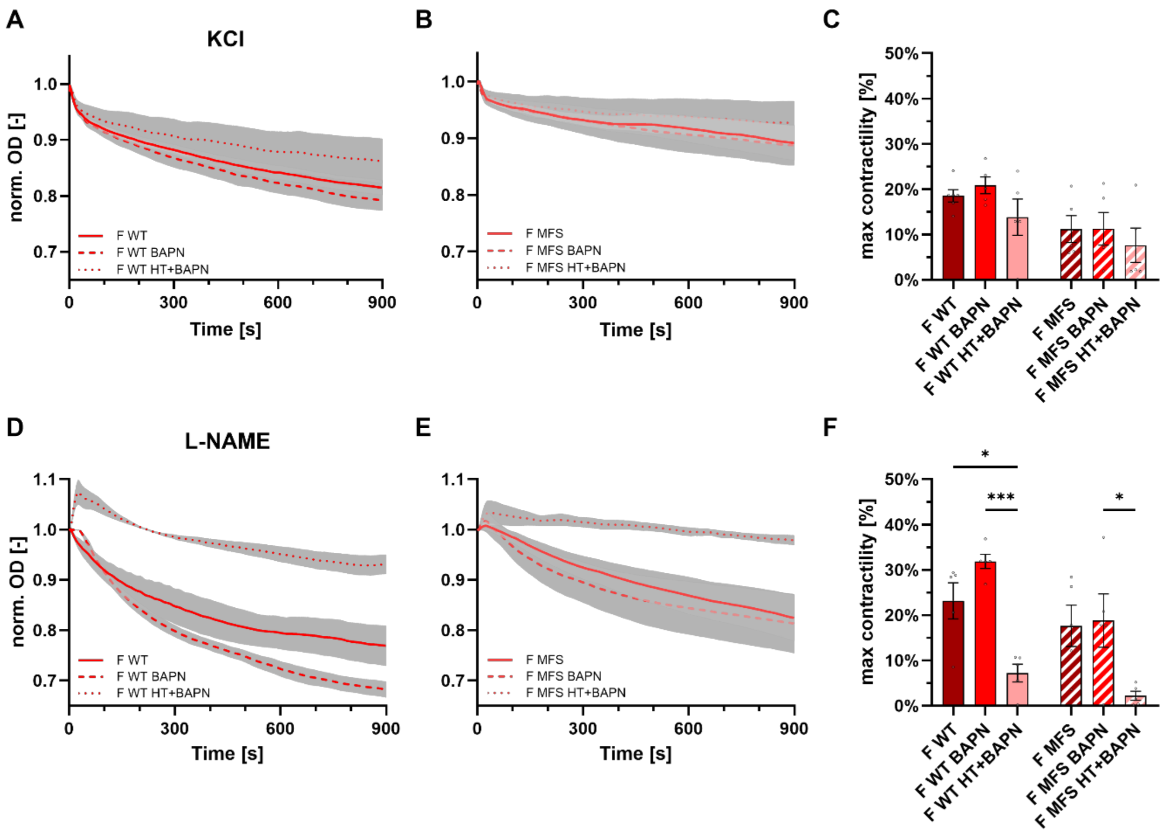


**Supplemental Figure S5**. **Vasoactive response to KCl and L-NAME showed similar loss in vasoactive function in female MFS mice.** Similar to Supplemental Figure S4 except for vasoactive response to the vasoconstrictors potassium chloride (KCl) and L-NAME, an eNOS inhibitor. Panels A, B, D, and E show time-course reductions in outer diameter in response to agonists while panels C and F show near steady-state maximum changes in outer diameter. Data expressed as mean ± SEM. Significance only shown within a genotype, indicated by * ($p$ < 0.05) and *** ($p$ < 0.0005).

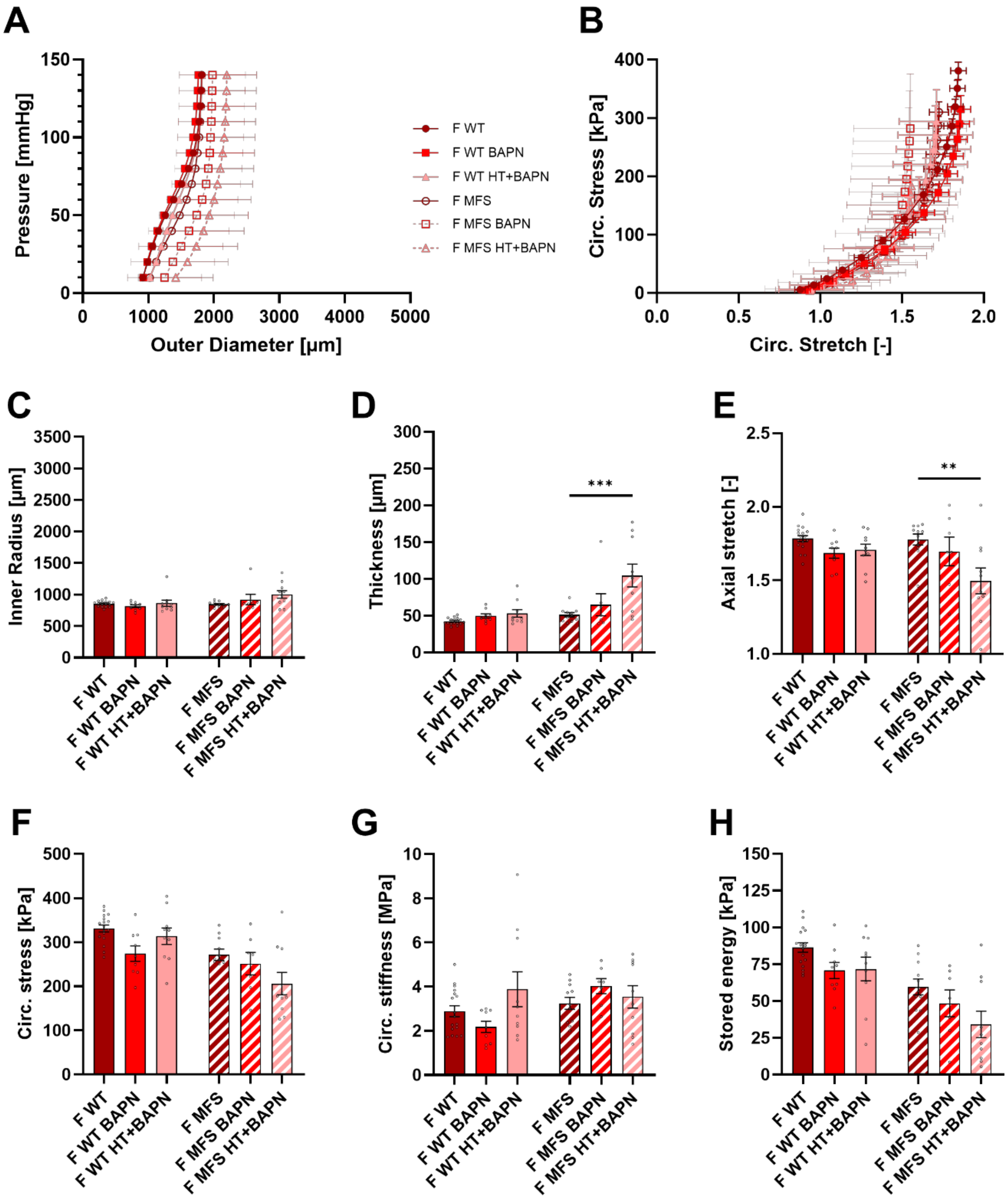


**Supplemental Figure S6. Female MFS mice experienced mild mechanical remodeling of the ascending aorta, with the greatest changes being wall thickening and loss of axial stretch in the HT+BAPN group.** Similar to Figure 3 in the main text, both observed and computed mechanical metrics for six female study groups: no-diet, BAPN, and combined HT+BAPN; *Fbn1*$^{+/+}$ (WT) and *Fbn1*$^{C1041G/+}$ (MFS). Panels A and B show overall pressure-diameter response and the computed circumferential stress-stretch relationship. Panels C-E show geometric remodeling while panels F-H show mechanical metrics computed from the four-fiber family constitutive equation. Data expressed as mean $\pm$ SEM. Significance only shown within a genotype, indicated by ** ($p < 0.005$) and *** ($p < 0.0005$). See Supplemental Figures S7-S8 for additional comparisons of passive mechanics.

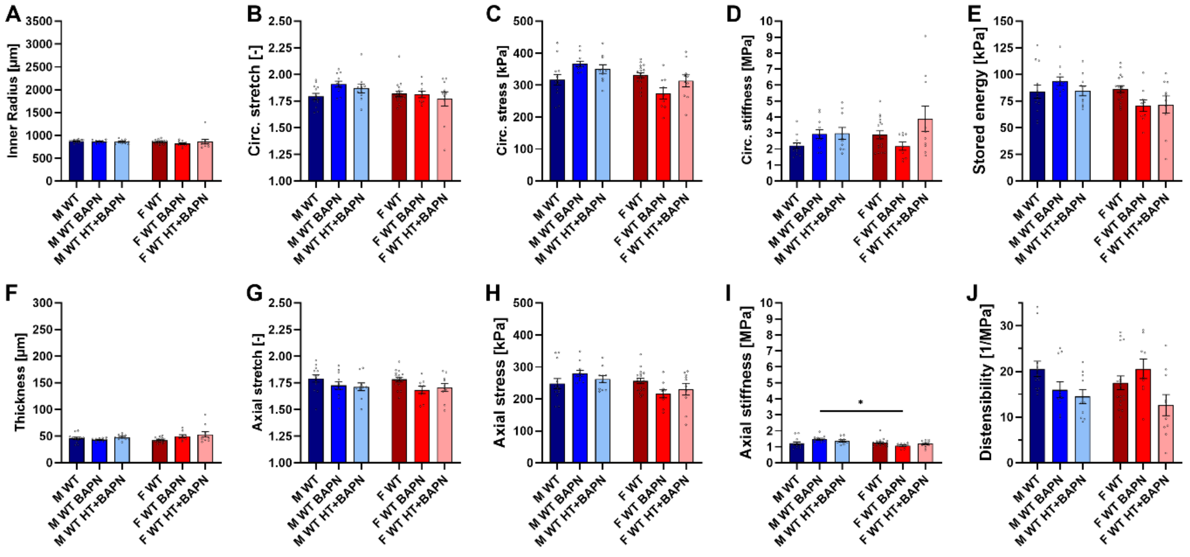


**Supplemental Figure S7. Additional passive mechanical metrics for all *Fbn1*$^{+/+}$ (WT) groups for examination of sex differences.** Data expressed as mean ± SEM. Significance only shown between males and females within a particular group for sex comparison purposes, indicated by * ($p < 0.05$).

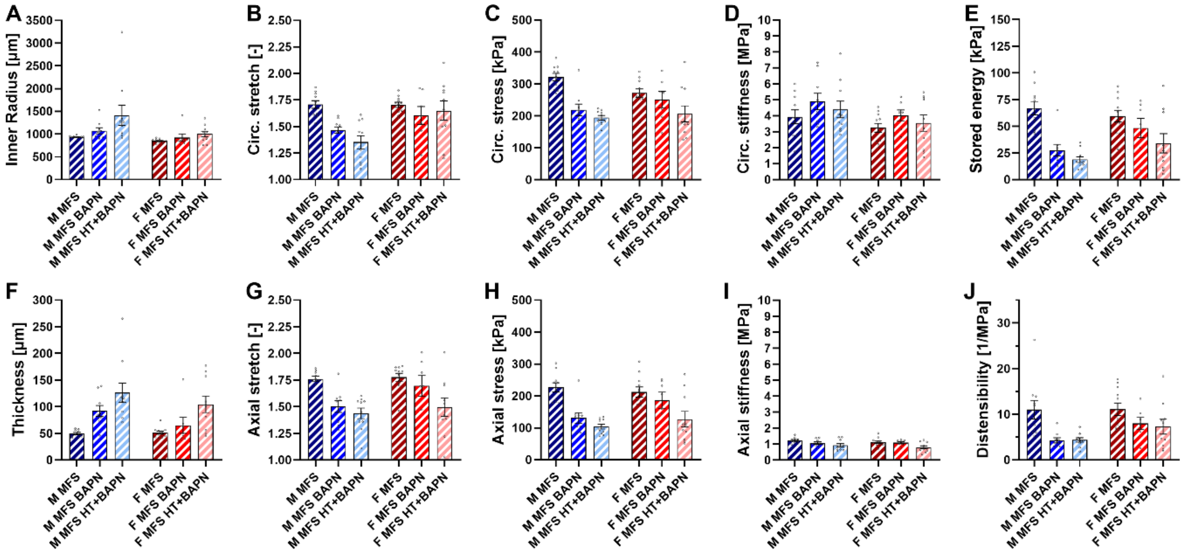


**Supplemental Figure S8. Additional passive mechanical metrics for all *Fbn1*$^{C1041G/+}$ (MFS) groups for examination of sex differences.** Data expressed as mean ± SEM. Significance only shown between males and females within a particular group for sex comparison purposes.

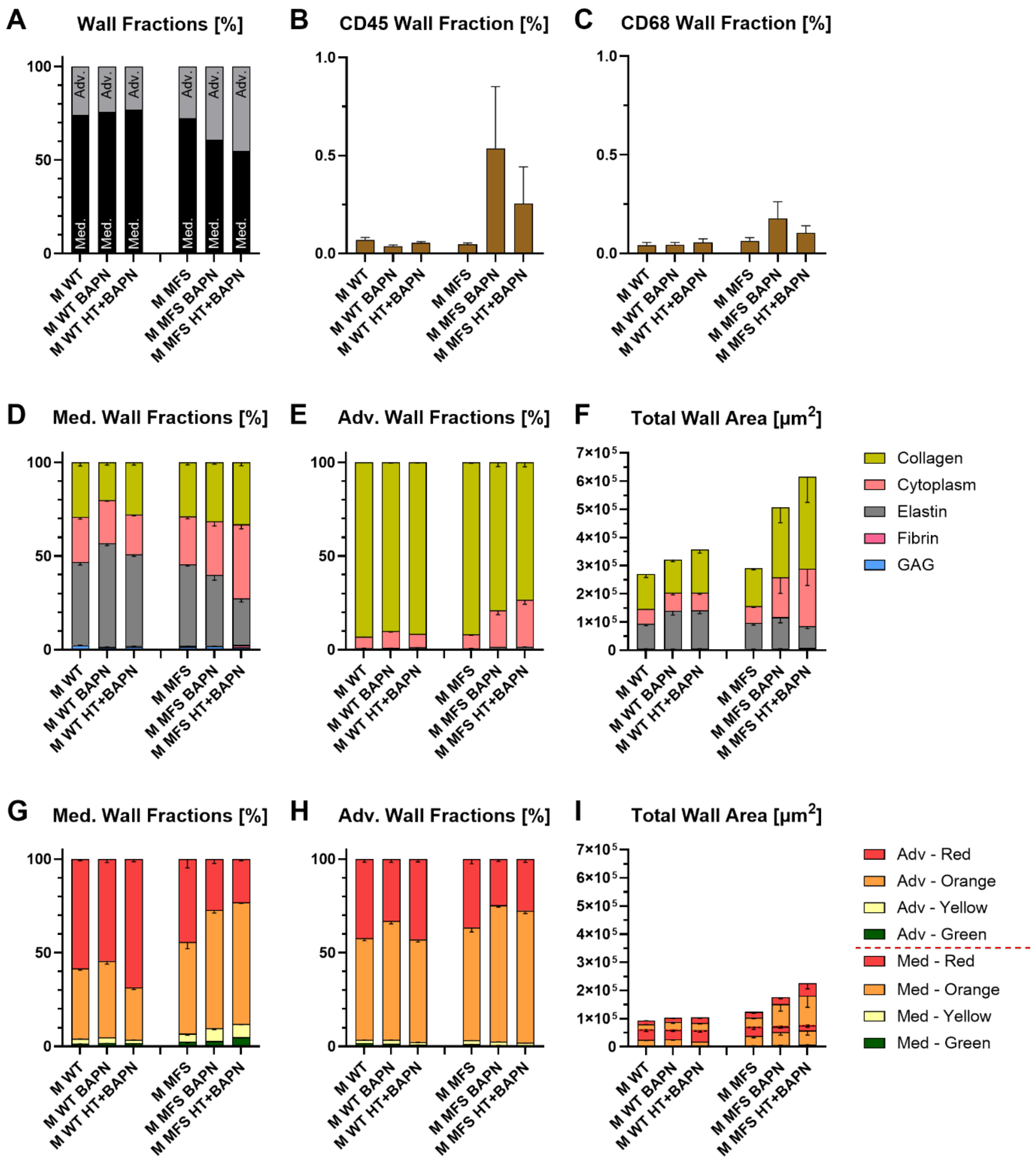


**Supplemental Figure S9. Combined Movat and PSR staining of unloaded aortic sections reveal dramatic adventitial thickening in male MFS groups in response to both BAPN and HT+BAPN.** Movat and PSR are used to establish medial vs. adventitial area fractions (A), demonstrating a shift in area in male MFS challenged by BAPN and HT+BAPN – two groups with marked increases in CD45 (B) and CD68 (C) positive areas. Panels D-F show quantification of extracellular constituents using Movat's Pentachrome while panels G-I show quantification of collagen fibers using picrosirius red. Data expressed as mean ± SEM. No statistics shown. See Supplemental Figure S10-S11 for similar data for female mice, which exhibited a milder phenotype.

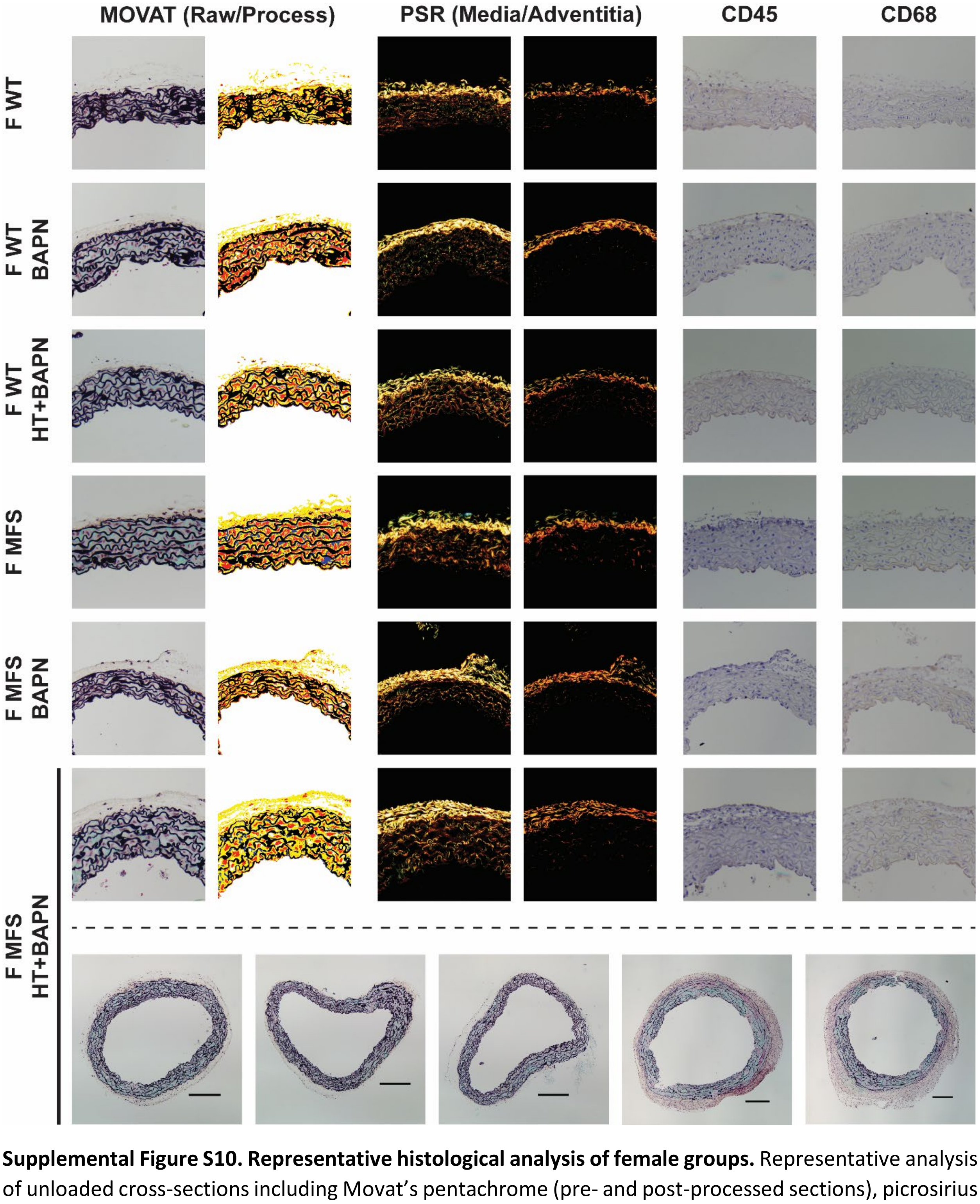


**Supplemental Figure S10. Representative histological analysis of female groups.** Representative analysis of unloaded cross-sections including Movat's pentachrome (pre- and post-processed sections), picrosirius red (captured at separate exposures to image specifically the media and adventitia), CD45, and CD68 staining. Female MFS HT+BAPN groups, with all five quantified samples shown in the bottom row, experienced a milder phenotype than corresponding male mice. Representative sectioned images are shown in a 250 µm x 250 µm box. Scale bars on complete sections of female MFS HT+BAPN samples (bottom row) indicate a 200 µm length. See Supplemental Figure S11 for quantifications of female mice.

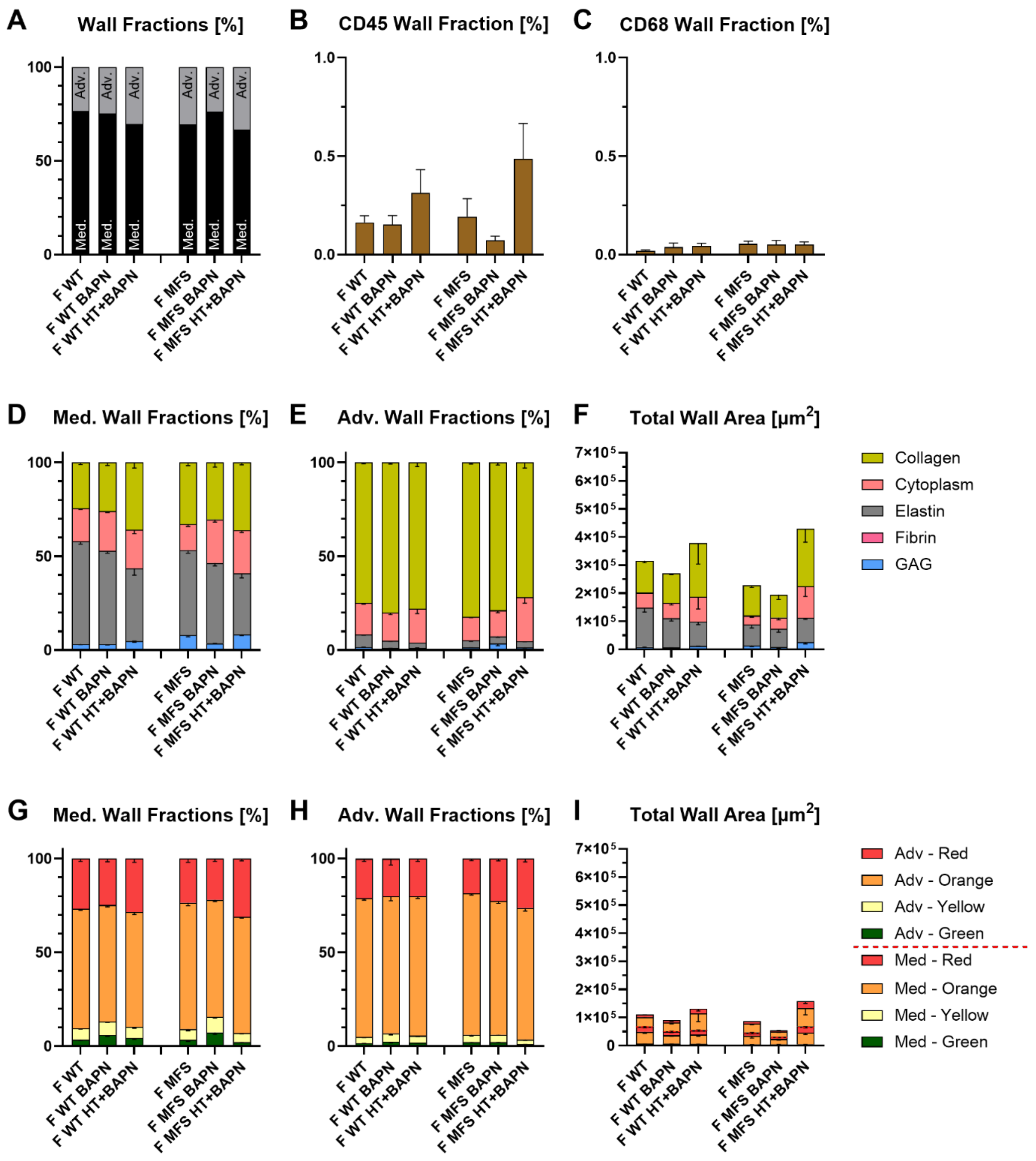


**Supplemental Figure S11. Combined Movat and PSR staining of unloaded aortic sections reveal reduced remodeling in female groups as compared to male.** Similar to Figure 4 in the main text, Movat and PSR are used to establish medial vs. adventitial area fractions (A) while immunohistochemical examination was performed to examine CD45 (B) and CD68 (C). Panels D-F show quantification of extracellular constituents using Movat's Pentachrome while panels G-I show quantification of collagen fibers using picrosirius red. Data expressed as mean ± SEM. No statistics shown.

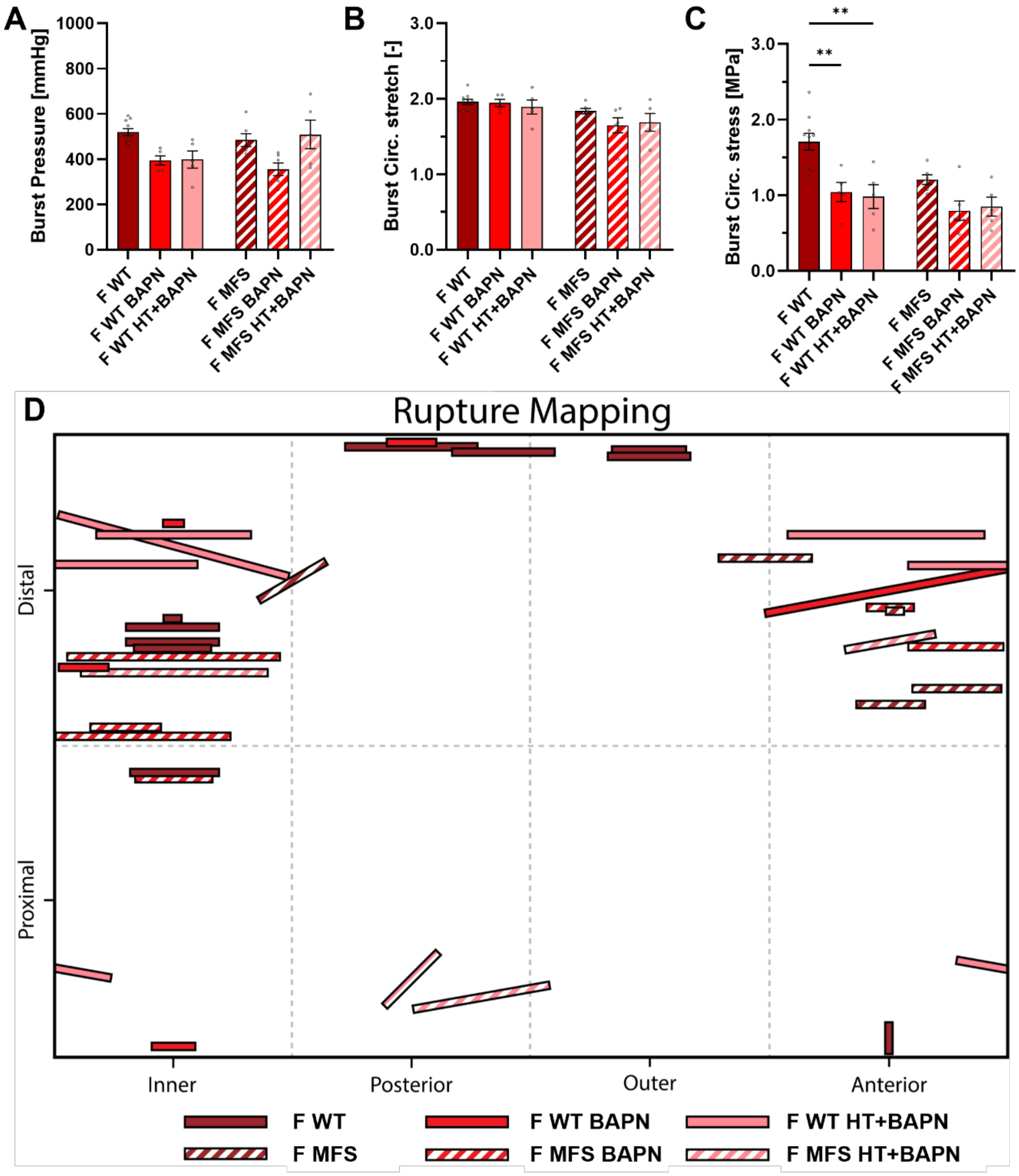


**Supplemental Figure 12. In female *Fbn1*$^{+/+}$ alone, BAPN and HT+BAPN diets induced a significant change in rupture properties.** Similar to Figure 7 in the main text, rupture mechanical analysis of six female study groups: no-diet, BAPN, and combined HT+BAPN; *Fbn1*$^{+/+}$ (WT) and *Fbn1*$^{C1041G/+}$ (MFS). Few significant changes occur in measured burst pressure (A), burst circumferential stretch (B), and computed circumferential stress at rupture (C). Panel D shows qualitatively the location, size, and orientation of rupture, noting a shift from distal regions to the aortic root in male MFS HT+BAPN mice. Data expressed as mean $\pm$ SEM. Significance only shown within a genotype.

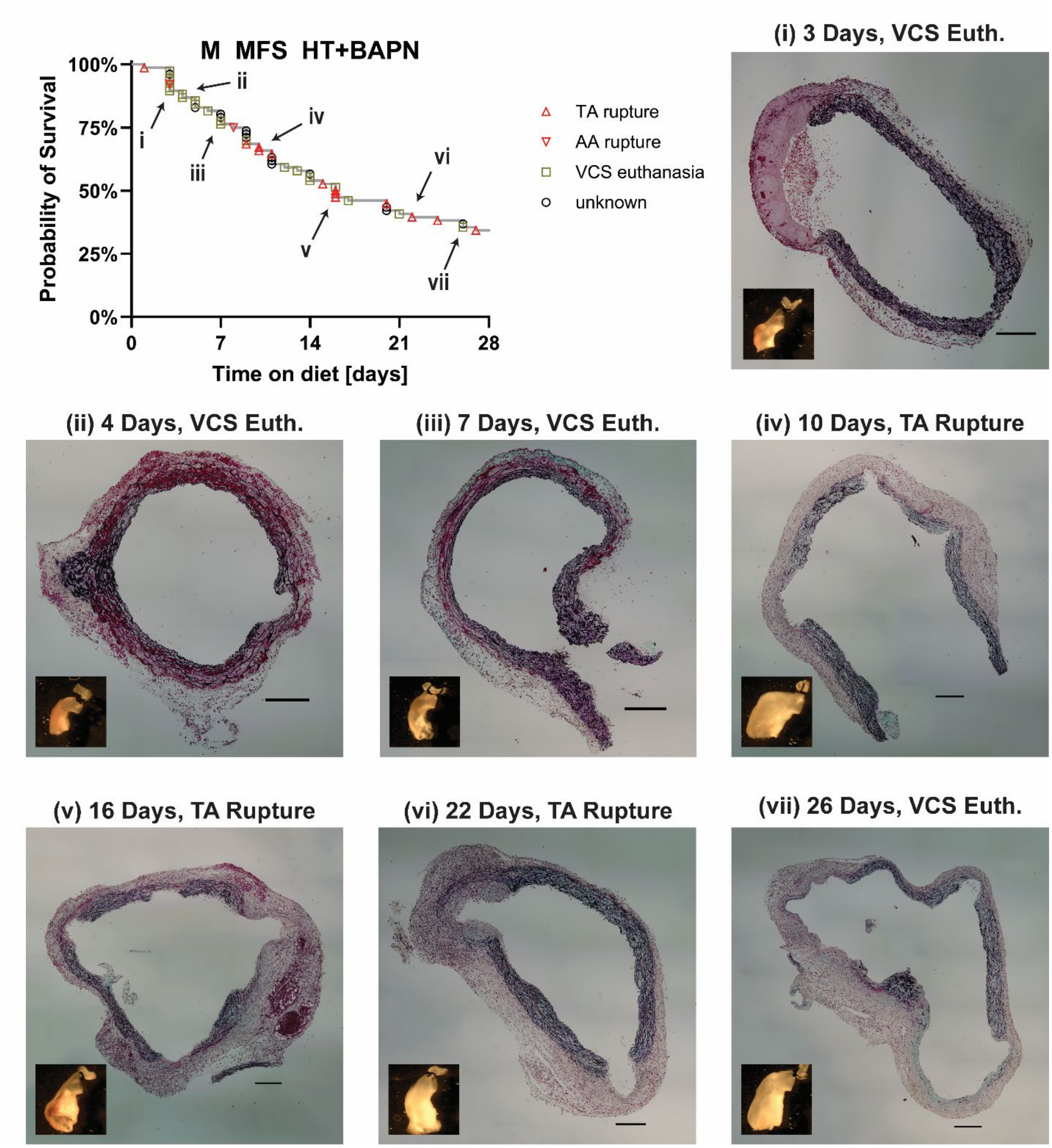


**Supplemental Figure S13.** Representative cross-sectional images of seven necropsied male MFS HT+BAPN samples, with duration of HT+BAPN and cause of death (VCS humane euthanasia or thoracic aortic rupture) as indicated. Scale bars on complete sections (black) indicate a 200 µm length. Gross vessel images are shown as 5 mm x 5mm insets.

**Supplemental Table 1.** Maximum normalized vasoactive responses through vasoactive experiments of all twelve groups: male (M) and female (F); *Fbn1*$^{+/+}$ (WT) and *Fbn1*$^{C1041G/+}$ (MFS); no-diet, BAPN diet, and combined HT+BAPN diet. Data expressed as mean $\pm$ standard deviation.

| | Ascending Thoracic Aorta - Males | | | | | |
|---|---|---|---|---|---|---|
| | **M WT** | **M WT BAPN** | **M WT HT+BAPN** | **M MFS** | **M MFS BAPN** | **M MFS HT+BAPN** |
| | **n = 5** | **n = 6** | **n = 5** | **n = 5** | **n = 5** | **n = 5** |
| **SMC max. response** | | | | | | |
| KCl (%) | 16.9% ± 8.3% | 17.3% ± 2.0% | 20.1% ± 3.7% | 9.8% ± 6.7% | 1.6% ± 2.2% | 0.9% ± 0.8% |
| AngII (%) | 7.7% ± 3.5% | 6.6% ± 5.3% | 10.4% ± 2.8% | 3.6% ± 2.2% | 3.3% ± 2.9% | 0.7% ± 0.7% |
| PE (%) | 30.1% ± 3.5% | 24.3% ± 7.9% | 29.1% ± 3.7% | 18.1% ± 7.2% | 5.9% ± 6.3% | 1.1% ± 1.3% |
| **EC max. response** | | | | | | |
| Ach (%) | 40.1% ± 10.6% | 35.2% ± 17.3% | 22.0% ± 7.4% | 23.8% ± 11.9% | 8.0% ± 7.5% | 0.3% ± 0.3% |
| L-NAME (%) | 28.0% ± 7.5% | 29.1% ± 4.9% | 15.1% ± 5.9% | 21.5% ± 4.5% | 5.3% ± 6.7% | 0.4% ± 0.2% |

| | Ascending Thoracic Aorta - Females | | | | | |
|---|---|---|---|---|---|---|
| | **F WT** | **F WT BAPN** | **F WT HT+BAPN** | **F MFS** | **F MFS BAPN** | **F MFS HT+BAPN** |
| | **n = 6** | **n = 5** | **n = 5** | **n = 5** | **n = 5** | **n = 5** |
| **SMC max. response** | | | | | | |
| KCl (%) | 18.5% ± 3.3% | 20.8% ± 4.2% | 13.8% ± 9.0% | 11.2% ± 6.6% | 11.2% ± 8.1% | 7.6% ± 8.5% |
| AngII (%) | 4.8% ± 2.8% | 9.4% ± 4.8% | 9.5% ± 5.2% | 4.5% ± 1.4% | 8.1% ± 5.1% | 9.6% ± 6.0% |
| PE (%) | 25.8% ± 4.1% | 30.7% ± 9.3% | 24.2% ± 13.3% | 19.0% ± 7.2% | 20.5% ± 13.2% | 16.7% ± 11.4% |
| **EC max. response** | | | | | | |
| Ach (%) | 29.9% ± 14.6% | 44.7% ± 16.2% | 10.5% ± 8.6% | 21.6% ± 11.5% | 27.5% ± 21.4% | 6.5% ± 6.5% |
| L-NAME (%) | 23.1% ± 8.9% | 31.9% ± 3.5% | 7.2% ± 4.4% | 17.6% ± 10.2% | 18.8% ± 13.1% | 2.2% ± 2.2% |

**Supplemental Table 2.** Group-averaged best-fit mechanical parameters to four-fiber family constitutive relationship. Maximum normalized vasoactive responses through vasoactive experiments of all twelve groups: male (M) and female (F); *Fbn1*$^{+/+}$ (WT) and *Fbn1*$^{C1041G/+}$ (MFS); no-diet, BAPN diet, and combined HT+BAPN diet.

| | | Elastic fibers | Axial Collagen | | Circ. Collagen +SMC | | Symmetric diagonal collagen | | | Error |
|---|---|---|---|---|---|---|---|---|---|---|
| | | c (kPa) | $c_1^1$ (kPa) | $c_2^1$ | $c_1^2$ (kPa) | $c_2^2$ | $c_1^{3,4}$ (kPa) | $c_2^{3,4}$ | $\alpha_o$ (deg) | *RMSE* |
| Ascending Thoracic Aorta | **M WT** | 18.963 | 3.956 | 0.189 | 4.202 | 1.183 | 9.432 | 0.233 | 49.218 | 0.084 |
| | **M WT BAPN** | 19.347 | 7.399 | 0.006 | 4.742 | 0.717 | 8.042 | 0.230 | 47.275 | 0.081 |
| | **M WT HT+BAPN** | 15.694 | 8.590 | 0.001 | 5.099 | 0.749 | 8.780 | 0.268 | 47.333 | 0.071 |
| | **M MFS** | 15.736 | 4.234 | 0.002 | 1.628 | 2.270 | 7.768 | 0.425 | 52.230 | 0.089 |
| | **M MFS BAPN** | 14.702 | 2.243 | 0.211 | 0.404 | 6.402 | 4.072 | 2.576 | 54.211 | 0.119 |
| | **M MFS HT+BAPN** | 14.879 | 0.876 | 0.839 | 0.377 | 2.636 | 15.436 | 6.193 | 57.499 | 0.153 |
| | **F WT** | 20.217 | 4.509 | 0.005 | 3.351 | 1.372 | 8.702 | 0.238 | 48.745 | 0.090 |
| | **F WT BAPN** | 16.119 | 7.060 | 0.003 | 3.040 | 1.164 | 9.331 | 0.263 | 47.581 | 0.085 |
| | **F WT HT+BAPN** | 14.057 | 7.015 | 0.005 | 2.748 | 4.117 | 9.208 | 0.984 | 50.505 | 0.093 |
| | **F MFS** | 13.262 | 4.756 | 0.007 | 0.003 | 2.359 | 7.402 | 0.385 | 52.777 | 0.091 |
| | **F MFS BAPN** | 12.237 | 4.919 | 0.002 | 0.024 | 2.440 | 7.673 | 2.416 | 53.201 | 0.105 |
| | **F MFS HT+BAPN** | 5.817 | 4.868 | 0.001 | 0.991 | 2.225 | 13.213 | 3.364 | 51.542 | 0.137 |

**Supplemental Table 3.** Key geometric and passive biomechanical metrics for six male (M) groups: *Fbn1*$^{+/+}$ (WT) and *Fbn1*$^{C1041G/+}$ (MFS); no-diet, BAPN diet, and combined HT+BAPN diet. Computed metrics are determined at group-averaged systolic blood pressures. Data expressed as mean ± standard deviation.

| | Ascending Thoracic Aorta - Males, Systolic BP | | | | | |
|---|---|---|---|---|---|---|
| | **M WT** | **M WT BAPN** | **M WT HT+BAPN** | **M MFS** | **M MFS BAPN** | **M MFS HT+BAPN** |
| | **n = 13** | **n = 13** | **n = 10** | **n = 10** | **n = 9** | **n = 10** |
| **Age (days)** | 84 ± 2 | 84 ± 1 | 84 ± 1 | 84 ± 1 | 84 ± 1 | 84 ± 2 |
| **Body mass (g)** | 24.71 ± 1.47 | 22.86 ± 2.28 | 19.76 ± 0.96 | 25.09 ± 1.20 | 25.56 ± 1.78 | 18.18 ± 1.90 |
| **Unloaded dimensions** | | | | | | |
| Wall Thickness (µm) | 146 ± 23 | 143 ± 9 | 152 ± 12 | 148 ± 10 | 195 ± 38 | 230 ± 59 |
| Outer Diameter (µm) | 1146 ± 75 | 1088 ± 75 | 1100 ± 89 | 1278 ± 108 | 1740 ± 413 | 2536 ± 1441 |
| Axial Length (mm) | 2.35 ± 0.19 | 2.30 ± 0.13 | 2.37 ± 0.16 | 2.76 ± 0.34 | 3.22 ± 0.31 | 3.62 ± 0.69 |
| **Loaded dimensions** | **P = 121** | **P = 137** | **P = 145** | **P = 126** | **P = 133** | **P = 130** |
| Outer Diameter (µm) | 1835 ± 72 | 1838 ± 102 | 1810 ± 89 | 1969 ± 72 | 2330 ± 446 | 3080 ± 1534 |
| Wall Thickness (µm) | 46 ± 8 | 43 ± 3 | 48 ± 5 | 50 ± 7 | 92 ± 30 | 126 ± 58 |
| Inner Radius (µm) | 872 ± 35 | 876 ± 51 | 857 ± 45 | 935 ± 33 | 1073 ± 197 | 1414 ± 710 |
| *in vivo* Axial Stretch ($\lambda_z^{iv}$) | 1.79 ± 0.12 | 1.76 ± 0.14 | 1.71 ± 0.11 | 1.76 ± 0.08 | 1.50 ± 0.17 | 1.44 ± 0.16 |
| *in vivo* Circumferential Stretch ($\lambda_\vartheta$) | 1.80 ± 0.10 | 1.90 ± 0.09 | 1.87 ± 0.13 | 1.71 ± 0.11 | 1.47 ± 0.09 | 1.35 ± 0.20 |
| **Systolic Cauchy Stresses (kPa)** | | | | | | |
| Circumferential, $\sigma_\vartheta$ | 317 ± 57.7 | 376 ± 32.7 | 350 ± 43.2 | 321 ± 37.6 | 219 ± 54.2 | 194 ± 21.9 |
| Axial, $\sigma_z$ | 248 ± 57.2 | 286 ± 30.9 | 262 ± 35.8 | 227 ± 45.3 | 131 ± 49.7 | 104 ± 23.4 |
| **Systolic Linearized Stiffness (MPa)** | | | | | | |
| Circumferential, $\mathscr{C}_{\theta\theta\theta\theta}$ | 2.21 ± 0.62 | 2.94 ± 0.88 | 2.98 ± 1.17 | 3.94 ± 1.38 | 4.89 ± 1.54 | 4.40 ± 1.65 |
| Axial, $\mathscr{C}_{zzzz}$ | 1.22 ± 0.32 | 1.48 ± 0.19 | 1.38 ± 0.22 | 1.23 ± 0.18 | 1.05 ± 0.23 | 0.93 ± 0.33 |
| **Systolic Stored Energy (kPa)** | 84 ± 22 | 97 ± 14 | 85 ± 15 | 67 ± 20 | 27 ± 16 | 19 ± 9 |
| **Distensibility (1/MPa)** | 20.55 ± 6.09 | 15.95 ± 5.38 | 14.47 ± 4.87 | 11.02 ± 6.39 | 4.29 ± 1.67 | 4.36 ± 1.50 |
| **PWV-MK (m/s)** | 7.48 ± 0.78 | 8.39 ± 1.29 | 8.93 ± 1.51 | 10.09 ± 2.20 | 14.09 ± 2.05 | 13.84 ± 2.44 |

**Supplemental Table 4.** Key geometric and passive biomechanical metrics for six female (F) groups: *Fbn1*$^{+/+}$ (WT) and *Fbn1*$^{C1041G/+}$ (MFS); no-diet, BAPN diet, and combined HT+BAPN diet. Computed metrics are determined at group-averaged systolic blood pressures. Data expressed as mean ± standard deviation.

| | Ascending Thoracic Aorta - Females, Systolic BP | | | | | |
|---|---|---|---|---|---|---|
| | **F WT** | **F WT BAPN** | **F WT HT+BAPN** | **F MFS** | **F MFS BAPN** | **F MFS HT+BAPN** |
| | **n = 18** | **n = 9** | **n = 10** | **n = 11** | **n = 8** | **n = 10** |
| **Age (days)** | 84 ± 1 | 83 ± 1 | 84 ± 1 | 84 ± 1 | 85 ± 2 | 84 ± 1 |
| **Body mass (g)** | 20.02 ± 1.97 | 18.42 ± 0.77 | 15.66 ± 1.73 | 20.61 ± 2.36 | 18.56 ± 0.70 | 15.55 ± 1.16 |
| **Unloaded dimensions** | | | | | | |
| Wall Thickness (µm) | 137 ± 9 | 151 ± 25 | 154 ± 17 | 153 ± 15 | 159 ± 34 | 231 ± 58 |
| Outer Diameter (µm) | 1099 ± 89 | 1080 ± 58 | 1193 ± 377 | 1181 ± 101 | 1370 ± 534 | 1590 ± 551 |
| Axial Length (mm) | 2.29 ± 0.19 | 2.31 ± 0.32 | 2.48 ± 0.45 | 2.71 ± 0.31 | 2.84 ± 0.45 | 2.94 ± 0.63 |
| **Loaded dimensions** | **P = 122** | **P = 122** | **P = 139** | **P = 120** | **P = 117** | **P = 138** |
| Outer Diameter (µm) | 1782 ± 108 | 1733 ± 127 | 1832 ± 339 | 1796 ± 92 | 1972 ± 514 | 2214 ± 466 |
| Wall Thickness (µm) | 42 ± 4 | 50 ± 8 | 53 ± 16 | 51 ± 10 | 65 ± 39 | 105 ± 49 |
| Inner Radius (µm) | 849 ± 53 | 817 ± 65 | 863 ± 157 | 847 ± 42 | 921 ± 219 | 1003 ± 189 |
| *in vivo* Axial Stretch ($\lambda_z^{iv}$) | 1.78 ± 0.08 | 1.68 ± 0.10 | 1.71 ± 0.12 | 1.78 ± 0.12 | 1.70 ± 0.26 | 1.50 ± 0.28 |
| *in vivo* Circumferential Stretch ($\lambda_\vartheta$) | 1.82 ± 0.11 | 1.81 ± 0.09 | 1.77 ± 0.21 | 1.70 ± 0.08 | 1.60 ± 0.22 | 1.65 ± 0.29 |
| **Systolic Cauchy Stresses (kPa)** | | | | | | |
| Circumferential, $\sigma_\vartheta$ | 329 ± 33.4 | 274 ± 52.0 | 314 ± 58.9 | 271 ± 42.5 | 251 ± 67.3 | 206 ± 80.5 |
| Axial, $\sigma_z$ | 257 ± 34.5 | 217 ± 39.5 | 230 ± 58.2 | 212 ± 53.4 | 187 ± 66.9 | 127 ± 77.4 |
| **Systolic Linearized Stiffness (MPa)** | | | | | | |
| Circumferential, $\mathscr{C}_{\theta\theta\theta\theta}$ | 2.80 ± 1.06 | 2.18 ± 0.76 | 3.89 ± 2.51 | 3.25 ± 0.90 | 4.03 ± 0.89 | 3.54 ± 1.60 |
| Axial, $\mathscr{C}_{zzzz}$ | 1.27 ± 0.24 | 1.07 ± 0.14 | 1.19 ± 0.21 | 1.13 ± 0.26 | 1.08 ± 0.18 | 0.79 ± 0.31 |
| **Systolic Stored Energy (kPa)** | 86 ± 13 | 71 ± 17 | 72 ± 25 | 60 ± 18 | 48 ± 24 | 34 ± 28 |
| **Distensibility (1/MPa)** | 18.20 ± 6.75 | 20.56 ± 6.42 | 12.61 ± 7.31 | 11.09 ± 4.51 | 7.99 ± 3.40 | 7.27 ± 5.16 |
| **PWV-MK (m/s)** | 8.19 ± 1.42 | 7.98 ± 1.49 | 10.43 ± 3.55 | 9.82 ± 1.81 | 11.46 ± 2.55 | 12.87 ± 4.15 |

**Supplemental Table 5.** Key geometric and passive biomechanical metrics for six male (M) groups: *Fbn1*$^{+/+}$ (WT) and *Fbn1*$^{C1041G/+}$ (MFS); no-diet, BAPN diet, and combined HT+BAPN diet. Computed metrics are determined at a blood pressure of 100 mmHg. Data expressed as mean $\pm$ standard deviation.

| | Ascending Thoracic Aorta - Males, Fixed BP | | | | | |
|---|---|---|---|---|---|---|
| | **M WT** | **M WT BAPN** | **M WT HT+BAPN** | **M MFS** | **M MFS BAPN** | **M MFS HT+BAPN** |
| | **n = 13** | **n = 13** | **n = 10** | **n = 10** | **n = 9** | **n = 10** |
| **Age (days)** | 84 ± 2 | 84 ± 1 | 84 ± 1 | 84 ± 1 | 84 ± 1 | 84 ± 2 |
| **Body mass (g)** | 24.71 ± 1.47 | 22.86 ± 2.28 | 19.76 ± 0.96 | 25.09 ± 1.20 | 25.56 ± 1.78 | 18.18 ± 1.90 |
| **Unloaded dimensions** | | | | | | |
| Wall Thickness (μm) | 146 ± 23 | 143 ± 9 | 152 ± 12 | 148 ± 10 | 195 ± 38 | 230 ± 59 |
| Outer Diameter (μm) | 1146 ± 75 | 1088 ± 75 | 1100 ± 89 | 1278 ± 108 | 1740 ± 413 | 2536 ± 1441 |
| Axial Length (mm) | 2.35 ± 0.19 | 2.30 ± 0.13 | 2.37 ± 0.16 | 2.76 ± 0.34 | 3.22 ± 0.31 | 3.62 ± 0.69 |
| **Loaded dimensions** | **P = 100** | **P = 100** | **P = 100** | **P = 100** | **P = 100** | **P = 100** |
| Outer Diameter (μm) | 1754 ± 86 | 1709 ± 102 | 1667 ± 114 | 1908 ± 100 | 2294 ± 448 | 3038 ± 1525 |
| Wall Thickness (μm) | 48 ± 8 | 46 ± 3 | 52 ± 5 | 51 ± 6 | 94 ± 30 | 128 ± 58 |
| Inner Radius (μm) | 829 ± 41 | 808 ± 51 | 781 ± 57 | 903 ± 47 | 1053 ± 198 | 1391 ± 705 |
| *in vivo* Axial Stretch ($\lambda_z^{iv}$) | 1.79 ± 0.12 | 1.76 ± 0.14 | 1.71 ± 0.11 | 1.76 ± 0.08 | 1.50 ± 0.17 | 1.44 ± 0.16 |
| *in vivo* Circumferential Stretch ($\lambda_\vartheta$) | 1.71 ± 0.09 | 1.76 ± 0.07 | 1.71 ± 0.09 | 1.65 ± 0.09 | 1.44 ± 0.08 | 1.33 ± 0.19 |
| **Fixed Cauchy Stresses (kPa)** | | | | | | |
| Circumferential, $\sigma_\vartheta$ | 237 ± 41.6 | 234 ± 23.3 | 201 ± 25.8 | 238 ± 24.1 | 158 ± 37.1 | 145 ± 15.9 |
| Axial, $\sigma_z$ | 214 ± 48.9 | 228 ± 23.2 | 200 ± 27.0 | 197 ± 35.1 | 110 ± 44.7 | 85 ± 19.6 |
| **Fixed Linearized Stiffness (MPa)** | | | | | | |
| Circumferential, $\mathscr{C}_{\theta\theta\theta\theta}$ | 1.35 ± 0.28 | 1.31 ± 0.26 | 1.14 ± 0.23 | 2.18 ± 0.62 | 2.69 ± 0.76 | 2.80 ± 1.22 |
| Axial, $\mathscr{C}_{zzzz}$ | 1.00 ± 0.25 | 1.09 ± 0.09 | 0.98 ± 0.14 | 1.02 ± 0.12 | 0.83 ± 0.19 | 0.71 ± 0.24 |
| **Fixed Stored Energy (kPa)** | 71 ± 18 | 74 ± 10 | 61 ± 9 | 57 ± 15 | 24 ± 13 | 16 ± 8 |
| **Distensibility (1/MPa)** | 20.55 ± 6.09 | 15.95 ± 5.38 | 14.47 ± 4.87 | 11.02 ± 6.39 | 4.29 ± 1.67 | 4.36 ± 1.50 |

**Supplemental Table 6.** Key geometric and passive biomechanical metrics for six female (F) groups: *Fbn1*$^{+/+}$ (WT) and *Fbn1*$^{C1041G/+}$ (MFS); no-diet, BAPN diet, and combined HT+BAPN diet. Computed metrics are determined at a blood pressure of 100 mmHg. Data expressed as mean ± standard deviation.

| | **Ascending Thoracic Aorta - Females, Fixed BP** | | | | | |
|---|---|---|---|---|---|---|
| | **F WT** | **F WT BAPN** | **F WT HT+BAPN** | **F MFS** | **F MFS BAPN** | **F MFS HT+BAPN** |
| | **n = 18** | **n = 9** | **n = 10** | **n = 11** | **n = 8** | **n = 10** |
| **Age (days)** | 84 ± 1 | 83 ± 1 | 84 ± 1 | 84 ± 1 | 85 ± 2 | 84 ± 1 |
| **Body mass (g)** | 20.02 ± 1.97 | 18.42 ± 0.77 | 15.66 ± 1.73 | 20.61 ± 2.36 | 18.56 ± 0.70 | 15.55 ± 1.16 |
| **Unloaded dimensions** | | | | | | |
| Wall Thickness (μm) | 137 ± 9 | 151 ± 25 | 154 ± 17 | 153 ± 15 | 159 ± 34 | 231 ± 58 |
| Outer Diameter (μm) | 1099 ± 89 | 1080 ± 58 | 1193 ± 377 | 1181 ± 101 | 1370 ± 534 | 1590 ± 551 |
| Axial Length (mm) | 2.29 ± 0.19 | 2.31 ± 0.32 | 2.48 ± 0.45 | 2.71 ± 0.31 | 2.84 ± 0.45 | 2.94 ± 0.63 |
| **Loaded dimensions** | **P = 100** | **P = 100** | **P = 100** | **P = 100** | **P = 100** | **P = 100** |
| Outer Diameter (μm) | 1708 ± 124 | 1658 ± 132 | 1727 ± 366 | 1755 ± 102 | 1947 ± 520 | 2149 ± 491 |
| Wall Thickness (μm) | 44 ± 4 | 52 ± 8 | 56 ± 16 | 52 ± 10 | 66 ± 39 | 108 ± 48 |
| Inner Radius (μm) | 810 ± 61 | 777 ± 67 | 807 ± 171 | 825 ± 46 | 908 ± 221 | 967 ± 201 |
| *in vivo* Axial Stretch ($\lambda_z^{iv}$) | 1.78 ± 0.08 | 1.68 ± 0.10 | 1.71 ± 0.12 | 1.78 ± 0.12 | 1.70 ± 0.26 | 1.50 ± 0.28 |
| *in vivo* Circumferential Stretch ($\lambda_\vartheta$) | 1.73 ± 0.09 | 1.73 ± 0.09 | 1.65 ± 0.16 | 1.66 ± 0.07 | 1.58 ± 0.21 | 1.58 ± 0.24 |
| **Fixed Cauchy Stresses (kPa)** | | | | | | |
| Circumferential, $\sigma_\vartheta$ | 245 ± 27.2 | 204 ± 38.6 | 196 ± 32.9 | 215 ± 30.6 | 208 ± 54.3 | 137 ± 49.0 |
| Axial, $\sigma_z$ | 224 ± 29.6 | 188 ± 34.7 | 186 ± 45.8 | 191 ± 44.3 | 174 ± 63.5 | 104 ± 66.0 |
| **Fixed Linearized Stiffness (MPa)** | | | | | | |
| Circumferential, $\mathcal{C}_{\theta\theta\theta\theta}$ | 1.49 ± 0.37 | 1.21 ± 0.27 | 1.63 ± 1.49 | 1.99 ± 0.46 | 2.66 ± 0.76 | 1.65 ± 0.73 |
| Axial, $\mathcal{C}_{zzzz}$ | 1.06 ± 0.19 | 0.90 ± 0.11 | 0.91 ± 0.15 | 1.00 ± 0.20 | 0.99 ± 0.19 | 0.61 ± 0.25 |
| **Fixed Stored Energy (kPa)** | 74 ± 9 | 59 ± 14 | 55 ± 18 | 53 ± 15 | 45 ± 22 | 27 ± 23 |
| **Distensibility (1/MPa)** | 18.20 ± 6.75 | 20.56 ± 6.42 | 12.61 ± 7.31 | 11.09 ± 4.51 | 7.99 ± 3.40 | 7.27 ± 5.16 |

**Supplemental Table 7.** (Immuno)histological quantification of unloaded sections stained using Movat's Pentachrome (MOVAT), Picrosirius Red (PSR), CD45, and CD68 for six male (M) groups: *Fbn1*$^{+/+}$ (WT) and *Fbn1*$^{C1041G/+}$ (MFS); no-diet, BAPN diet, and combined HT+BAPN diet. Data expressed as mean $\pm$ standard deviation.

| | Ascending Thoracic Aorta - Male Histology | | | | | |
|---|---|---|---|---|---|---|
| | M WT | M WT BAPN | M WT HT+BAPN | M MFS | M MFS BAPN | M MFS HT+BAPN |
| **Total Wall Fractions** | | | | | | |
| Media [%] | 74.0% ± 3.9% | 75.7% ± 3.4% | 76.9% ± 3.9% | 72.3% ± 3.5% | 60.8% ± 15.8% | 54.9% ± 18.2% |
| Adventitia [%] | 26.0% ± 3.9% | 24.3% ± 3.4% | 23.1% ± 3.9% | 27.7% ± 3.5% | 39.2% ± 15.8% | 45.1% ± 18.2% |
| **Total Wall Areas** | | | | | | |
| Media [µm^2] | 199,513 ± 21,296 | 242,391 ± 42,061 | 273,250 ± 32,617 | 210,221 ± 20,352 | 282,279 ± 57,267 | 292,671 ± 73,248 |
| Adventitia [µm^2] | 70,177 ± 14,801 | 79,625 ± 23,595 | 83,378 ± 24,213 | 80,013 ± 8,214 | 224,810 ± 194,508 | 322,757 ± 267,012 |
| **MOVAT** | | | | | | |
| **Media Area Fractions** | | | | | | |
| GAG [%] | 2.3% ± 0.7% | 1.2% ± 0.4% | 1.7% ± 0.5% | 1.5% ± 0.6% | 1.7% ± 0.7% | 1.3% ± 0.6% |
| Fibrin [%] | 0.1% ± 0.3% | 0.3% ± 0.3% | 0.3% ± 0.3% | 0.6% ± 0.8% | 0.3% ± 0.4% | 1.2% ± 1.6% |
| Elastin [%] | 44.2% ± 6.5% | 55.3% ± 5.7% | 49.0% ± 5.2% | 43.4% ± 3.4% | 37.9% ± 12.2% | 24.8% ± 8.0% |
| Cytoplasm [%] | 24.2% ± 5.0% | 22.9% ± 2.7% | 21.2% ± 3.6% | 25.6% ± 5.2% | 28.7% ± 11.1% | 39.6% ± 10.4% |
| Collagen [%] | 29.1% ± 8.6% | 20.2% ± 6.5% | 27.8% ± 6.5% | 28.8% ± 5.2% | 31.5% ± 4.4% | 33.1% ± 7.7% |
| **Adventitia Area Fractions** | | | | | | |
| GAG [%] | 0.4% ± 0.2% | 0.3% ± 0.1% | 0.4% ± 0.2% | 0.2% ± 0.1% | 0.3% ± 0.1% | 0.2% ± 0.1% |
| Fibrin [%] | 0.0% ± 0.0% | 0.0% ± 0.0% | 0.1% ± 0.1% | 0.0% ± 0.0% | 0.0% ± 0.0% | 0.1% ± 0.1% |
| Elastin [%] | 0.4% ± 0.1% | 0.6% ± 0.1% | 0.7% ± 0.2% | 0.5% ± 0.2% | 1.1% ± 0.7% | 1.3% ± 0.7% |
| Cytoplasm [%] | 6.1% ± 1.1% | 9.1% ± 2.2% | 7.3% ± 1.4% | 7.4% ± 1.9% | 19.5% ± 9.9% | 25.0% ± 10.2% |
| Collagen [%] | 93.1% ± 1.1% | 90.0% ± 2.2% | 91.5% ± 1.5% | 91.8% ± 2.0% | 79.0% ± 10.2% | 73.4% ± 10.4% |
| **PSR** | | | | | | |
| **Collagen Wall Area** | | | | | | |
| Media [µm^2] | 60,622 ± 22,345 | 58,195 ± 15,812 | 57,803 ± 15,633 | 60,123 ± 13,600 | 71,043 ± 32,271 | 74,932 ± 52,671 |
| Adventitia [µm^2] | 32,908 ± 4,019 | 45,324 ± 17,698 | 46,667 ± 16,778 | 46,348 ± 8,985 | 105,295 ± 71,638 | 150,226 ± 136,135 |
| **Media Area Fractions** | | | | | | |
| Red [%] | 58.5% ± 3.6% | 54.4% ± 7.3% | 68.6% ± 5.6% | 44.3% ± 18.4% | 27.2% ± 9.5% | 23.0% ± 4.0% |
| Orange [%] | 37.4% ± 3.2% | 40.7% ± 6.6% | 27.8% ± 3.9% | 49.0% ± 14.0% | 63.1% ± 6.2% | 64.9% ± 2.5% |
| Yellow [%] | 2.5% ± 0.5% | 3.0% ± 0.9% | 2.0% ± 0.8% | 4.3% ± 2.8% | 6.8% ± 2.6% | 7.1% ± 1.9% |
| Green [%] | 1.6% ± 0.8% | 1.8% ± 0.9% | 1.6% ± 1.1% | 2.5% ± 2.1% | 2.9% ± 1.6% | 5.0% ± 2.2% |
| **Adventitia Area Fractions** | | | | | | |
| Red [%] | 42.2% ± 6.6% | 33.1% ± 7.1% | 43.0% ± 6.1% | 36.5% ± 10.0% | 24.7% ± 4.8% | 27.6% ± 7.0% |
| Orange [%] | 54.1% ± 5.3% | 63.3% ± 5.8% | 54.7% ± 5.1% | 60.2% ± 9.1% | 72.8% ± 4.0% | 70.4% ± 5.7% |
| Yellow [%] | 2.0% ± 0.8% | 2.3% ± 0.8% | 1.4% ± 0.7% | 2.1% ± 0.8% | 1.8% ± 0.7% | 1.4% ± 1.0% |
| Green [%] | 1.7% ± 0.9% | 1.4% ± 0.6% | 0.9% ± 0.6% | 1.1% ± 0.4% | 0.7% ± 0.3% | 0.5% ± 0.4% |
| **IHC** | | | | | | |
| CD45 area fraction [%] | 0.07% ± 0.02% | 0.04% ± 0.01% | 0.06% ± 0.02% | 0.05% ± 0.02% | 0.54% ± 0.70% | 0.25% ± 0.42% |
| CD68 area fraction [%] | 0.04% ± 0.03% | 0.04% ± 0.03% | 0.06% ± 0.04% | 0.06% ± 0.04% | 0.18% ± 0.19% | 0.10% ± 0.08% |

**Supplemental Table 8.** (Immuno)histological quantification of unloaded sections stained using Movat's Pentachrome (MOVAT), Picrosirius Red (PSR), CD45, and CD68 for six female (F) groups: *Fbn1*$^{+/+}$ (WT) and *Fbn1*$^{C1041G/+}$ (MFS); no-diet, BAPN diet, and combined HT+BAPN diet. Data expressed as mean $\pm$ standard deviation.

| | Ascending Thoracic Aorta - Female Histology | | | | | |
|---|---|---|---|---|---|---|
| | F WT | F WT BAPN | F WT HT+BAPN | F MFS | F MFS BAPN | F MFS HT+BAPN |
| **Total Wall Fractions** | | | | | | |
| Media [%] | 76.6% ± 1.9% | 75.4% ± 0.9% | 69.7% ± 10.4% | 69.3% ± 12.8% | 76.2% ± 6.8% | 66.5% ± 10.3% |
| Adventitia [%] | 23.4% ± 1.9% | 24.6% ± 0.9% | 30.3% ± 10.4% | 30.7% ± 12.8% | 23.8% ± 6.8% | 33.5% ± 10.3% |
| **Total Wall Areas** | | | | | | |
| Media [µm^2] | 242,545 ± 44,078 | 203,710 ± 27,811 | 244,196 ± 113,158 | 162,380 ± 57,306 | 151,685 ± 66,116 | 268,581 ± 80,878 |
| Adventitia [µm^2] | 73,335 ± 8,129 | 66,656 ± 10,354 | 134,530 ± 149,848 | 65,225 ± 11,382 | 42,910 ± 10,845 | 160,502 ± 127,428 |
| **MOVAT** | | | | | | |
| **Media Area Fractions** | | | | | | |
| GAG [%] | 3.3% ± 1.1% | 3.1% ± 1.2% | 4.8% ± 2.2% | 8.0% ± 3.1% | 3.6% ± 1.4% | 8.4% ± 2.4% |
| Fibrin [%] | 0.0% ± 0.0% | 0.0% ± 0.0% | 0.1% ± 0.1% | 0.0% ± 0.0% | 0.0% ± 0.1% | 0.0% ± 0.0% |
| Elastin [%] | 54.7% ± 5.8% | 49.7% ± 4.3% | 38.8% ± 13.7% | 45.3% ± 5.8% | 42.8% ± 6.1% | 32.4% ± 9.4% |
| Cytoplasm [%] | 17.5% ± 2.3% | 21.3% ± 2.8% | 20.5% ± 7.4% | 13.9% ± 4.4% | 23.2% ± 5.3% | 23.0% ± 4.2% |
| Collagen [%] | 24.5% ± 4.7% | 25.9% ± 5.6% | 35.9% ± 11.0% | 32.8% ± 5.9% | 30.4% ± 9.9% | 36.2% ± 4.9% |
| **Adventitia Area Fractions** | | | | | | |
| GAG [%] | 1.8% ± 0.5% | 1.1% ± 0.6% | 1.2% ± 0.6% | 1.5% ± 0.8% | 3.5% ± 4.7% | 1.6% ± 1.8% |
| Fibrin [%] | 0.0% ± 0.0% | 0.0% ± 0.0% | 0.0% ± 0.1% | 0.0% ± 0.0% | 0.0% ± 0.0% | 0.0% ± 0.0% |
| Elastin [%] | 6.4% ± 1.1% | 3.9% ± 1.3% | 2.8% ± 1.2% | 3.7% ± 1.6% | 3.8% ± 1.9% | 3.0% ± 0.6% |
| Cytoplasm [%] | 17.0% ± 2.5% | 14.9% ± 3.4% | 18.0% ± 9.3% | 12.5% ± 1.6% | 13.8% ± 5.0% | 23.6% ± 12.3% |
| Collagen [%] | 74.8% ± 2.8% | 80.0% ± 2.8% | 78.0% ± 8.0% | 82.3% ± 2.8% | 78.8% ± 5.2% | 71.8% ± 11.6% |
| **PSR** | | | | | | |
| **Collagen Wall Area** | | | | | | |
| Media [µm^2] | 65,931 ± 15,454 | 48,776 ± 11,741 | 53,326 ± 18,021 | 44,178 ± 21,867 | 30,078 ± 12,684 | 65,947 ± 20,450 |
| Adventitia [µm^2] | 44,726 ± 7,176 | 41,196 ± 11,416 | 77,861 ± 84,271 | 42,892 ± 14,292 | 24,924 ± 6,221 | 92,085 ± 71,685 |
| **Media Area Fractions** | | | | | | |
| Red [%] | 26.7% ± 5.6% | 24.8% ± 6.0% | 28.6% ± 8.2% | 23.6% ± 7.1% | 22.1% ± 5.9% | 31.1% ± 5.2% |
| Orange [%] | 63.7% ± 3.5% | 62.1% ± 3.5% | 61.1% ± 4.6% | 67.3% ± 5.6% | 62.3% ± 3.6% | 61.9% ± 2.7% |
| Yellow [%] | 6.1% ± 1.9% | 7.2% ± 1.3% | 6.0% ± 2.6% | 5.8% ± 3.0% | 8.3% ± 2.5% | 4.8% ± 1.8% |
| Green [%] | 3.5% ± 2.0% | 5.9% ± 2.5% | 4.3% ± 2.6% | 3.3% ± 2.9% | 7.3% ± 3.8% | 2.2% ± 1.1% |
| **Adventitia Area Fractions** | | | | | | |
| Red [%] | 21.0% ± 5.5% | 20.1% ± 11.6% | 20.2% ± 5.7% | 18.7% ± 4.1% | 22.7% ± 5.0% | 26.5% ± 7.4% |
| Orange [%] | 74.0% ± 4.4% | 73.0% ± 8.1% | 74.2% ± 4.3% | 75.4% ± 3.3% | 71.3% ± 4.5% | 70.1% ± 6.0% |
| Yellow [%] | 3.3% ± 1.0% | 4.5% ± 2.6% | 3.7% ± 1.8% | 3.8% ± 1.6% | 3.8% ± 1.0% | 2.4% ± 1.6% |
| Green [%] | 1.6% ± 0.6% | 2.3% ± 1.6% | 1.9% ± 1.2% | 2.2% ± 1.2% | 2.2% ± 0.8% | 1.1% ± 0.9% |
| **IHC** | | | | | | |
| CD45 area fraction [%] | 0.16% ± 0.08% | 0.15% ± 0.09% | 0.31% ± 0.26% | 0.19% ± 0.18% | 0.07% ± 0.05% | 0.48% ± 0.40% |
| CD68 area fraction [%] | 0.02% ± 0.01% | 0.04% ± 0.04% | 0.04% ± 0.03% | 0.05% ± 0.02% | 0.05% ± 0.05% | 0.05% ± 0.03% |

**Supplemental Table 9.** Statistical comparisons of histological and IHC quantifications between six male groups: *Fbn1*$^{+/+}$ (WT) and *Fbn1*$^{C1041G/+}$ (MFS); no-diet, BAPN diet, and combined HT+BAPN diet. Statistical differences are indicated by n.s. (not significant), * ($p < 0.05$), ** ($p < 0.005$), *** ($p < 0.0005$), and **** ($p < 0.0001$).

| Ascending Thoracic Aorta - Male Histology, Statistics | M WT vs M WT BAPN | M WT vs M WT HT+BAPN | M WT vs M MFS | M WT vs M MFS BAPN | M WT vs M MFS HT+BAPN | M WT BAPN vs M WT HT+BAPN | M WT BAPN vs M MFS | M WT BAPN vs M MFS BAPN | M WT BAPN vs M MFS HT+BAPN | M WT HT+BAPN vs M MFS | M WT HT+BAPN vs M MFS BAPN | M WT HT+BAPN vs M MFS HT+BAPN | M MFS vs M MFS BAPN | M MFS vs M MFS HT+BAPN | M MFS BAPN vs M MFS HT+BAPN |
|---|---|---|---|---|---|---|---|---|---|---|---|---|---|---|---|
| **MOVAT** | | | | | | | | | | | | | | | |
| **Media Area Fractions** | | | | | | | | | | | | | | | |
| GAG [%] | **** | n.s. | ** | n.s. | **** | n.s. | n.s. | n.s. | n.s. | n.s. | n.s. | n.s. | n.s. | n.s. | n.s. |
| Fibrin [%] | * | n.s. | n.s. | n.s. | * | n.s. | n.s. | n.s. | n.s. | n.s. | n.s. | n.s. | n.s. | n.s. | n.s. |
| Elastin [%] | ** | n.s. | n.s. | n.s. | *** | n.s. | ** | **** | **** | n.s. | n.s. | **** | n.s. | * | n.s. |
| Cytoplasm [%] | n.s. | n.s. | n.s. | n.s. | *** | n.s. | n.s. | n.s. | **** | n.s. | * | **** | n.s. | n.s. | n.s. |
| Collagen [%] | * | n.s. | n.s. | n.s. | n.s. | n.s. | * | *** | **** | n.s. | n.s. | n.s. | n.s. | n.s. | n.s. |
| **Adventitia Area Fractions** | | | | | | | | | | | | | | | |
| GAG [%] | * | n.s. | * | n.s. | ** | n.s. | n.s. | n.s. | n.s. | n.s. | n.s. | * | n.s. | n.s. | n.s. |
| Fibrin [%] | * | * | n.s. | n.s. | *** | n.s. | n.s. | n.s. | n.s. | n.s. | n.s. | n.s. | n.s. | * | n.s. |
| Elastin [%] | n.s. | *** | n.s. | **** | **** | n.s. | n.s. | n.s. | ** | n.s. | n.s. | n.s. | * | *** | n.s. |
| Cytoplasm [%] | * | n.s. | n.s. | **** | **** | n.s. | n.s. | n.s. | ** | n.s. | ** | **** | ** | **** | n.s. |
| Collagen [%] | * | n.s. | n.s. | **** | **** | n.s. | n.s. | n.s. | ** | n.s. | ** | **** | ** | **** | n.s. |
| **PSR** | | | | | | | | | | | | | | | |
| **Media Area Fractions** | | | | | | | | | | | | | | | |
| Red [%] | n.s. | n.s. | n.s. | ** | *** | n.s. | n.s. | ** | ** | ** | **** | **** | n.s. | n.s. | n.s. |
| Orange [%] | n.s. | n.s. | n.s. | *** | *** | n.s. | n.s. | ** | ** | ** | **** | **** | n.s. | n.s. | n.s. |
| Yellow [%] | n.s. | n.s. | n.s. | ** | *** | n.s. | n.s. | * | ** | n.s. | **** | **** | n.s. | * | n.s. |
| Green [%] | n.s. | n.s. | n.s. | n.s. | *** | n.s. | n.s. | n.s. | ** | n.s. | n.s. | *** | n.s. | * | n.s. |
| **Adventitia Area Fractions** | | | | | | | | | | | | | | | |
| Red [%] | n.s. | n.s. | n.s. | *** | ** | n.s. | n.s. | n.s. | n.s. | n.s. | **** | *** | ** | n.s. | n.s. |
| Orange [%] | n.s. | n.s. | n.s. | **** | **** | n.s. | n.s. | n.s. | n.s. | n.s. | **** | **** | ** | n.s. | n.s. |
| Yellow [%] | n.s. | n.s. | n.s. | n.s. | n.s. | * | n.s. | n.s. | * | n.s. | n.s. | n.s. | n.s. | n.s. | n.s. |
| Green [%] | n.s. | n.s. | n.s. | * | **** | n.s. | n.s. | n.s. | *** | n.s. | n.s. | n.s. | n.s. | ** | n.s. |
| **IHC** | | | | | | | | | | | | | | | |
| CD45 area fraction [%] | n.s. | n.s. | n.s. | * | n.s. | n.s. | n.s. | **** | n.s. | n.s. | ** | n.s. | **** | n.s. | n.s. |
| CD68 area fraction [%] | n.s. | n.s. | n.s. | ** | n.s. | n.s. | n.s. | * | n.s. | n.s. | n.s. | n.s. | n.s. | n.s. | n.s. |

**Supplemental Table 10.** Statistical comparisons of histological and IHC quantifications between six female groups: *Fbn1*$^{+/+}$ (WT) and *Fbn1*$^{C1041G/+}$ (MFS); no-diet, BAPN diet, and combined HT+BAPN diet. Statistical differences are indicated by n.s. (not significant), * ($p < 0.05$), ** ($p < 0.005$), *** ($p < 0.0005$), and **** ($p < 0.0001$).

**Ascending Thoracic Aorta - Female Histology, Statistics**

| | F WT vs F WT BAPN | F WT vs F WT HT+BAPN | F WT vs F MFS | F WT vs F MFS BAPN | F WT vs F MFS HT+BAPN | F WT BAPN vs F WT HT+BAPN | F WT BAPN vs F MFS | F WT BAPN vs F MFS BAPN | F WT BAPN vs F MFS HT+BAPN | F WT HT+BAPN vs F MFS | F WT HT+BAPN vs F MFS BAPN | F WT HT+BAPN vs F MFS HT+BAPN | F MFS vs F MFS BAPN | F MFS vs F MFS HT+BAPN | F MFS BAPN vs F MFS HT+BAPN |
|---|---|---|---|---|---|---|---|---|---|---|---|---|---|---|---|
| **MOVAT** | | | | | | | | | | | | | | | |
| **Media Area Fractions** | | | | | | | | | | | | | | | |
| GAG [%] | n.s. | n.s. | *** | n.s. | **** | n.s. | *** | n.s. | **** | n.s. | n.s. | * | ** | n.s. | **** |
| Fibrin [%] | n.s. | n.s. | ** | n.s. | n.s. | n.s. | * | n.s. | n.s. | n.s. | n.s. | n.s. | * | n.s. | n.s. |
| Elastin [%] | n.s. | *** | n.s. | ** | **** | n.s. | n.s. | n.s. | *** | n.s. | n.s. | n.s. | n.s. | n.s. | n.s. |
| Cytoplasm [%] | n.s. | n.s. | n.s. | * | * | n.s. | ** | n.s. | n.s. | n.s. | n.s. | n.s. | *** | *** | n.s. |
| Collagen [%] | n.s. | ** | n.s. | n.s. | **** | n.s. | n.s. | n.s. | ** | n.s. | n.s. | n.s. | n.s. | n.s. | n.s. |
| **Adventitia Area Fractions** | | | | | | | | | | | | | | | |
| GAG [%] | n.s. | n.s. | n.s. | n.s. | n.s. | n.s. | n.s. | n.s. | n.s. | n.s. | n.s. | n.s. | n.s. | n.s. | n.s. |
| Fibrin [%] | n.s. | n.s. | n.s. | n.s. | n.s. | n.s. | *** | n.s. | * | n.s. | n.s. | n.s. | n.s. | n.s. | n.s. |
| Elastin [%] | * | **** | * | ** | **** | n.s. | n.s. | n.s. | n.s. | n.s. | n.s. | n.s. | n.s. | n.s. | n.s. |
| Cytoplasm [%] | n.s. | n.s. | * | n.s. | n.s. | n.s. | n.s. | n.s. | n.s. | n.s. | n.s. | n.s. | n.s. | * | n.s. |
| Collagen [%] | n.s. | n.s. | *** | n.s. | n.s. | n.s. | n.s. | n.s. | n.s. | n.s. | n.s. | n.s. | n.s. | n.s. | n.s. |
| **PSR** | | | | | | | | | | | | | | | |
| **Media Area Fractions** | | | | | | | | | | | | | | | |
| Red [%] | n.s. | n.s. | n.s. | n.s. | n.s. | n.s. | n.s. | n.s. | n.s. | n.s. | n.s. | n.s. | n.s. | n.s. | ** |
| Orange [%] | n.s. | n.s. | n.s. | n.s. | n.s. | n.s. | n.s. | n.s. | n.s. | n.s. | n.s. | n.s. | n.s. | n.s. | n.s. |
| Yellow [%] | n.s. | n.s. | n.s. | n.s. | n.s. | n.s. | n.s. | n.s. | n.s. | n.s. | n.s. | n.s. | n.s. | n.s. | ** |
| Green [%] | n.s. | n.s. | n.s. | n.s. | n.s. | n.s. | n.s. | n.s. | ** | n.s. | n.s. | n.s. | * | n.s. | *** |
| **Adventitia Area Fractions** | | | | | | | | | | | | | | | |
| Red [%] | n.s. | n.s. | n.s. | n.s. | n.s. | n.s. | n.s. | n.s. | * | n.s. | n.s. | n.s. | n.s. | n.s. | n.s. |
| Orange [%] | n.s. | n.s. | n.s. | n.s. | n.s. | n.s. | n.s. | n.s. | n.s. | n.s. | n.s. | n.s. | n.s. | n.s. | n.s. |
| Yellow [%] | n.s. | n.s. | n.s. | n.s. | n.s. | n.s. | n.s. | n.s. | * | n.s. | n.s. | n.s. | n.s. | n.s. | n.s. |
| Green [%] | n.s. | n.s. | n.s. | n.s. | n.s. | n.s. | n.s. | n.s. | n.s. | n.s. | n.s. | n.s. | n.s. | n.s. | * |
| **IHC** | | | | | | | | | | | | | | | |
| CD45 area fraction [%] | n.s. | n.s. | n.s. | n.s. | n.s. | n.s. | n.s. | n.s. | n.s. | n.s. | * | n.s. | n.s. | n.s. | *** |
| CD68 area fraction [%] | n.s. | n.s. | n.s. | n.s. | * | n.s. | n.s. | n.s. | n.s. | n.s. | n.s. | n.s. | n.s. | n.s. | n.s. |

**Supplemental Table 11.** Quantification of rupture mechanics of all twelve groups: male (M) and female (F); *Fbn1*$^{+/+}$ (WT) and *Fbn1*$^{C1041G/+}$ (MFS); no-diet, BAPN diet, and combined HT+BAPN diet. Data expressed as mean $\pm$ standard deviation.

| | Ascending Thoracic Aorta - Males | | | | | |
|---|---|---|---|---|---|---|
| | **M WT** | **M WT BAPN** | **M WT HT+BAPN** | **M MFS** | **M MFS BAPN** | **M MFS HT+BAPN** |
| | **n = 7** | **n = 6** | **n = 5** | **n = 5** | **n = 5** | **n = 5** |
| **Measured Metrics** | | | | | | |
| Burst Pressure (mmHg) | 509 ± 44 | 383 ± 65 | 373 ± 125 | 509 ± 48 | 435 ± 50 | 462 ± 225 |
| Burst OD (µm) | 2079 ± 68 | 2000 ± 127 | 1921 ± 87 | 2189 ± 26 | 2529 ± 250 | 2899 ± 1140 |
| **Calculated Metrics** | | | | | | |
| Loaded inner radius (µm) | 996 ± 40 | 961 ± 63 | 914 ± 44 | 1043 ± 11 | 1175 ± 100 | 1291 ± 437 |
| Loaded Thickness (µm) | 44 ± 6 | 40 ± 2 | 46 ± 3 | 51 ± 4 | 90 ± 27 | 158 ± 141 |
| Circumferential Stretch (-) | 2.02 ± 0.14 | 2.05 ± 0.07 | 1.92 ± 0.08 | 1.76 ± 0.09 | 1.63 ± 0.11 | 1.39 ± 0.38 |
| Circumferential Stress (kPa) | 1568.4 ± 266.1 | 1246.0 ± 255.9 | 983.7 ± 327.3 | 1391.4 ± 100.1 | 790.1 ± 179.3 | 745.0 ± 538.7 |
| Circ. Stress "Factor of Safety" (vs sys) | 5.19 ± 0.69 | 3.37 ± 0.65 | 2.89 ± 1.20 | 4.78 ± 0.48 | 3.96 ± 0.57 | 3.84 ± 2.42 |

| | Ascending Thoracic Aorta - Females | | | | | |
|---|---|---|---|---|---|---|
| | **F WT** | **F WT BAPN** | **F WT HT+BAPN** | **F MFS** | **F MFS BAPN** | **F MFS HT+BAPN** |
| | **n = 10** | **n = 5** | **n = 5** | **n = 6** | **n = 6** | **n = 5** |
| **Burst Measured** | | | | | | |
| Burst Pressure (mmHg) | 519 ± 50 | 394 ± 46 | 398 ± 84 | 484 ± 70 | 355 ± 68 | 509 ± 142 |
| Burst OD (µm) | 2027 ± 156 | 1906 ± 73 | 1889 ± 152 | 2042 ± 112 | 2142 ± 451 | 2401 ± 474 |
| **Calculated Metrics** | | | | | | |
| Loaded inner radius (µm) | 973 ± 80 | 905 ± 45 | 894 ± 81 | 969 ± 53 | 1002 ± 183 | 1101 ± 185 |
| Loaded Thickness (µm) | 40 ± 5 | 48 ± 9 | 51 ± 12 | 52 ± 7 | 69 ± 44 | 100 ± 53 |
| Circumferential Stretch (-) | 1.96 ± 0.11 | 1.95 ± 0.11 | 1.89 ± 0.21 | 1.84 ± 0.09 | 1.65 ± 0.24 | 1.69 ± 0.26 |
| Circumferential Stress (kPa) | 1706.4 ± 348.9 | 1041.6 ± 284.1 | 981.4 ± 350.3 | 1206.2 ± 159.7 | 794.8 ± 312.8 | 850.5 ± 279.1 |
| Circ. Stress "Factor of Safety" (vs sys) | 5.11 ± 0.75 | 3.76 ± 0.51 | 3.20 ± 0.72 | 5.00 ± 1.09 | 3.47 ± 0.78 | 4.31 ± 1.50 |
